# UBE2G1 Governs the Destruction of Cereblon Neomorphic Substrates

**DOI:** 10.1101/389098

**Authors:** Gang Lu, Stephanie Weng, Mary Matyskiela, Xinde Zheng, Wei Fang, Scott Wood, Christine Surka, Reina Mizukoshi, Chin-Chun Lu, Derek Mendy, In Sock Jang, Kai Wang, Mathieu Marella, Suzana Couto, Brian Cathers, James Carmichael, Philip Chamberlain, Mark Rolfe

## Abstract

The immunomodulatory drugs (IMiDs) thalidomide, lenalidomide, and pomalidomide as well as the novel cereblon modulating agents (CMs) including CC-122, CC-220 and cereblon-based proteolysis-targeting chimaeras (PROTACs) repurpose the Cul4-RBX1-DDB1-CRBN (CRL4^CRBN^) E3 ubiquitin ligase complex to induce the degradation of specific neomorphic substrates via polyubiquitination in conjunction with an E1 ubiquitin-activating enzyme and E2 ubiquitin-conjugating enzymes, which have until now remained elusive. Here we show that the ubiquitin-conjugating enzymes UBE2G1 and UBE2D3 cooperatively promote the polyubiquitination of CRL4^CRBN^ neomorphic substrates in a cereblon- and CM-dependent manner via a sequential ubiquitination mechanism: UBE2D3 transforms the neomorphic substrates into mono-ubiquitinated forms, upon which UBE2G1 catalyzes K48-linked polyubiquitin chain extension. Blockade of UBE2G1 diminishes the ubiquitination and degradation of neomorphic substrates, and consequent antitumor activities elicited by all tested CMs. For example, UBE2G1 inactivation significantly attenuated the degradation of myeloma survival factors IKZF1 and IKZF3 induced by lenalidomide and pomalidomide, hence conferring drug resistance. UBE2G1-deficient myeloma cells, however, remained sensitive to a more potent IKZF1/3 degrader CC-220. Collectively, these findings suggest that loss of UBE2G1 activity might be a resistance mechanism to drugs that hijack the CRL4^CRBN^ to eliminate disease-driving proteins, and that this resistance mechanism can be overcome by next-generation CMs that destroy the same targeted protein more effectively.

## Introduction

The ubiquitin-proteasome system (UPS) is a highly regulated component of the protein homeostasis network that dictates multiple cellular processes in eukaryotes (Hershko and Ciechanover, 1998). Through the orchestrated actions of ubiquitin-activating enzymes (E1), ubiquitin-conjugating enzymes (E2) and ubiquitin-ligating enzymes (E3), the ε-amine of a lysine residue in a target protein is covalently conjugated with K48- or K11-linked poly-ubiquitin chains, thereby marking the target protein for proteasomal degradation (Jin et al., 2008; Komander and Rape, 2012; Pickart, 2001). Recently, repurposing the Cullin-Ring E3 ligase complexes CRL4^CRBN^ (Cul4-RBX1-DDB1-CRBN) and CRL2^VHL^ (Cul2-RBX1-EloB/C-VHL) with small-molecule degraders to remove disease-driving proteins otherwise considered ‘undruggable’ has emerged as a novel therapeutic modality that has the potential to transform drug discovery and development (Bondeson and Crews, 2017; Huang and Dixit, 2016; Lebraud and Heightman, 2017).

There are two types of small-molecule degraders that have been exploited used to date. The first is represented by thalidomide (THAL), lenalidomide (LEN) and pomalidome (POM), as well as other cereblon modulating agents CC-122, CC-220, and CC-885. This class of molecule docks into a tri-tryptophan pocket in the thalidomide-binding domain of cereblon, the substrate receptor of CRL4^CRBN^, to create a hotspot for protein-protein interactions thereby enhancing the binding of unique neomorphic substrate to cereblon, resulting in substrate ubiquitination and degradation (Fischer et al., 2014) (Chamberlain et al., 2014) (Matyskiela et al., 2018) (Petzold et al., 2016). THAL, LEN and POM promote the degradation of two hematopoietic transcription factors IKZF1 and IKZF3 to achieve anti-myeloma activity (Kronke et al., 2014) (Lu et al., 2014a) (Gandhi et al., 2014), whereas only LEN targets CK1α for effective degradation (Kronke et al., 2015), which is presumably linked to its efficacy in myelodysplastic syndrome with chromosome 5q deletion. CC-220 is a significantly more potent IKZF1 and IKZF3 degrader than IMiD drugs (Matyskiela et al., 2018) (Nakayama et al., 2017) (Schafer et al., 2018), and it is currently in clinical trials for relapsed/refractory multiple myeloma and systemic lupus erythematosus. By contrast, CC-885 is the only aforementioned cereblon modulating agent that allows cereblon to recognize translation termination factor GSPT1 for ubiquitination and degradation (Matyskiela et al., 2016). The second type of small-molecule degraders is generally referred to as a proteolysis-targeting chimera (PROTAC) (Sakamoto et al., 2001), which is composed of two linked protein binding ligands, with one engaging a target protein and the other interacting with an E3 ubiquitin ligase such as CRL4^CRBN^ or CRL2^VHL^ to trigger proximity-induced substrate ubiquitination and degradation (Deshaies, 2015) (Neklesa et al., 2017). Many PROTACs have been described recently, but the clinical value of this approach has not yet been established.

CRL4^CRBN^ belongs to the multi-subunit cullin-RING E3 ubiquitin ligase family containing over 200 members (Petroski and Deshaies, 2005). The mammalian cullin scaffold proteins (including Cul1, Cul2, Cul3, Cul4A, Cul4B, Cul5, and Cul7) bring their substrates into close proximity with E2 ubiquitin-conjugating enzymes, thereby enabling effective substrate ubiquitination (Petroski and Deshaies, 2005). SCF(Skp1-Cul1-F-box)^Cdc4^, the founding member of the cullin-RING E3 ligase family, was first discovered in the budding yeast *Saccharomyces cerevisiae*, in which SCF^Cdc4^ works in conjunction with a single E2 ubiquitin-conjugating enzyme Cdc34 to promote the polyubiquitination of a variety of SCF substrates (Feldman et al., 1997) (Skowyra et al., 1997) (Blondel et al., 2000) (Jang et al., 2001; Perkins et al., 2001). Human Cdc34/UBE2R1 can substitute for yeast Cdc34 in *Saccharomyces cerevisiae* underscoring their functional conservation (Plon et al., 1993). However, in contrast to its dominant role in catalyzing the ubiquitination of SCF substrates in yeast, Cdc34 coordinates ubiquitination with UBE2D3/UbcH5c via a sequential ubiquitination mechanism to improve reaction rate and efficiency in human cells. In brief, Cdc34 acts as an ubiquitin chain elongation enzyme that assembles the K48-linked ubiquitin chains on mono-ubiquitins pre-conjugated to SCF substrates by UBE2D3 (Pan et al., 2004). Such sequential ubiquitination by two E2 enzymes was first reported for the anaphase-promoting complex ubiquitin ligase (Rodrigo-Brenni and Morgan, 2007). Several ubiquitin conjugation E2 enzymes have been reported to regulate CRL4 substrates as well. For instance, in response to UV irradiation, the CRL4^cdt2^ ligase complex mediates the proteolysis of Cdt1 with the help of E2 enzymes UBE2G1 and its paralog UBE2G2, while working together with a different E2 enzyme UbcH8/UBEL6 to trigger the degradation of p21 and Set8 in human cells (Shibata et al., 2011). Despite the proven cellular efficacy and clinical success of many cereblon modulating agents, it remain unknown whether unique ubiquitin E2 enzymes control the ubiquitination of each specific cereblon neomorphic substrate, and whether loss of E2 enzymes contributes to resistance to these agents.

## Results

### UBE2G1 is the dominant ubiquitin E2 enzyme that governs the destruction of cereblon neomorphic substrates induced by cereblon modulating agents

The clinical course of multiple myeloma typically follows a recurring pattern of remission and relapse with resistance to IMiD drugs based combination regimens (Harousseau and Attal, 2017). Such relapse is not frequently associated with cereblon downregulation and/or mutation (Kortum et al., 2016; Qian et al., 2018) (Zhu et al., 2011)Hence, we reasoned that resistance to IMiD drugs in myeloma could be ascribed to reduced degradation of IKZF1 and IKZF3 as a result of inactivation of other essential components of the CRL4^CRBN^ ligase complex, for instance the E2 ubiquitin conjugation enzyme. To look for such proteins, we devised a high-throughput CRISPR-Cas9 screen approach to monitor the effect of individual knockout of a gene of interest on POM-induced degradation of IKZF1 protein tagged with enhanced ProLabel (ePL), a small β-galactosidase N-terminal fragment (Figure 1A), and created a single guide RNA (sgRNA) library containing three sgRNAs for each of the 41 annotated E2 enzymes in the human genome, as well as three non-targeting sgRNAs in arrayed format (Supplemental Table 1). The ePL tag complements with the large β-galactosidase C-terminal fragment to form an active enzyme that hydrolyzes substrate to emit a chemiluminescent signal, allowing the measurement of ePL-IKZF1 fusion protein level in a high-throughput fashion.

**Figure 1.**
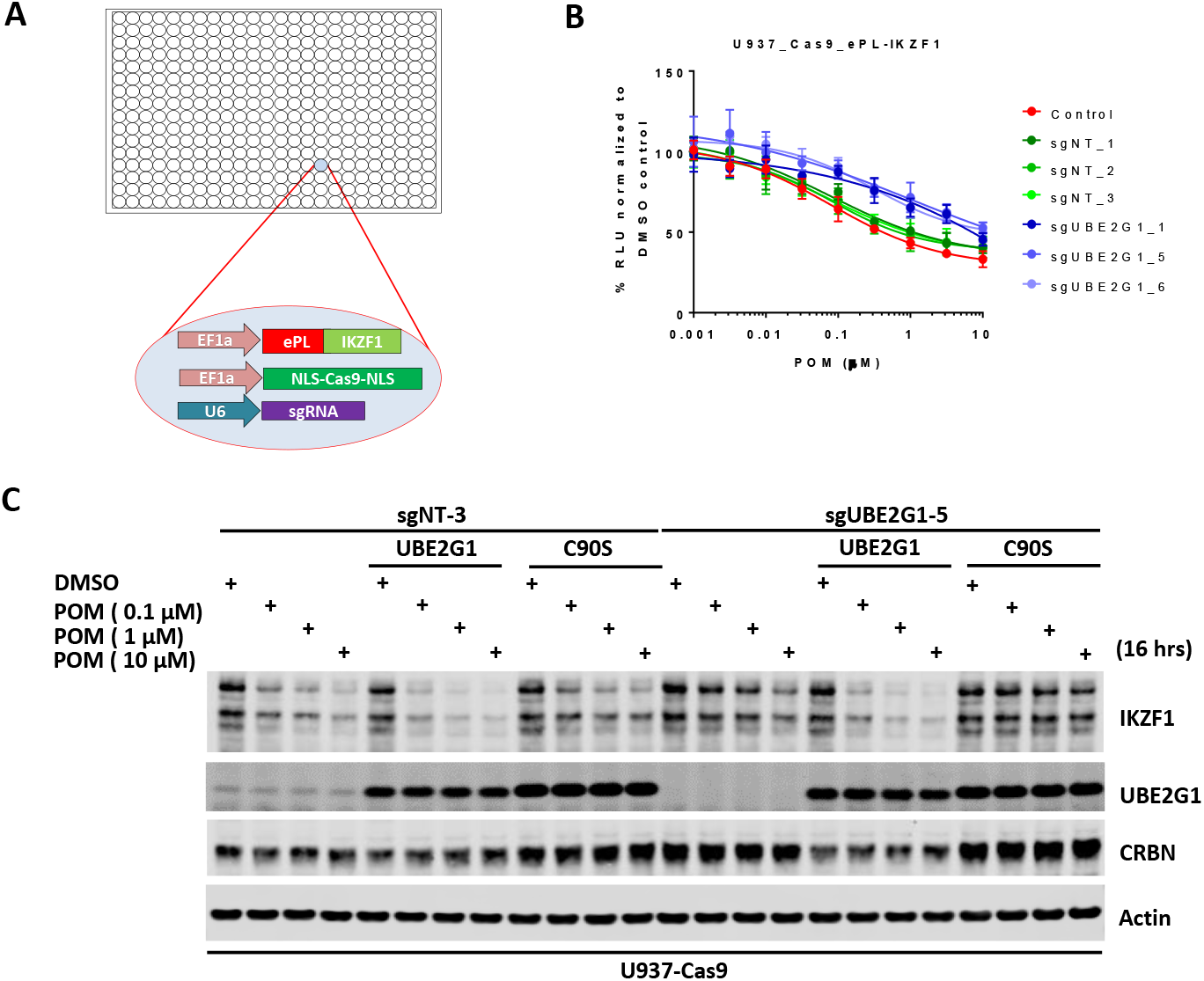
Identification of UBE2G1 as the most critical ubiquitin E2 enzyme that mediates the pomalidomide-induced degradation of IKZF1. (A) Schematic showing the design of the CRISPR screen to identify E2 enzyme(s) regulating the CM-induced degradation of ePL-tagged IKZF1 in 384-well array format. (B) Chemiluminescent measurement of ePL-IKZF1 protein expression level in U937-Cas9_ePL-IKZF1 parental cells or cells expressing non-targeting or UBE2G1-specific sgRNAs. Cells were treated with POM at the indicated concentrations for 16 hours. Data are presented as mean ± SD (n = 4). (C) Immunoblot analysis of U937-Cas9 parental or UBE2G1-/- cells with or without stable expression of UBE2G1 wild-type or C90S mutant. Cells were treated with POM at the indicated concentrations for 16 hours.

To determine the robustness of this screening approach, we transduced U937 cells stably expressing Cas9 and ePL-IKZF1 (U937_Cas9_ePL-IKZF1) with lentiviral vector expressing a non-targeting or *CRBN*-specific sgRNA. Four days post transduction, cells were seeded into 384-well plates pre-dispensed with either DMSO vehicle control or POM at varying concentrations. Sixteen hours after incubation, IKZF1 degradation was assessed using the ePL luminescent assay. As expected, cereblon knockout completely abrogated the degradation of ePL-tagged IKZF1 fusion protein (Figure S1A). Using this approach we then evaluated the effect of individual knockout of each E2 enzyme on ePL-IKZF1 degradation induced by POM. Out of 41 E2 enzymes, UBE2G1 and to a lesser extent UBE2M, UBE2D3, and UBE2D2/UbcH5b, when depleted, imposed statistically significant inhibition on the ePL-IKZF1 degradation (Figures 1B, S2D and S2I).

UBE2M, also called UBC12, is a NEDD8-conjuating enzyme, which regulates, via neddylation, the activity of all Cullin Ring E3 ligases including CRL4^CRBN^ (Petroski and Deshaies, 2005) (Gong and Yeh, 1999) (Pan et al., 2004). Indeed, co-treatment with MLN4924, an inhibitor of NEDD8-activating enzyme (Soucy et al., 2009), prevented the degradation of ePL-IKZF1 induced by POM (Figure S1C). The effect of UBE2M knockout, however, was much less pronounced (Figure S2I). Given the well-established role of UBE2M in cell proliferation and survival, we reasoned that this difference could be explained by that U937 cells only with partial UBE2M loss survived four days after CRISPR gene editing. Consistent with this notion, cellular fitness was markedly reduced by 48-hour treatment of MLN2924 at concentrations that elicited near-complete blockage of POM-induced IKZF1 degradation in the U937_Cas9_ePL-IKZF1 cell line used in the screen (Figures S1C and S1D).

CRISPR knockout of UBE2G1 also attenuated the destabilization of endogenous IKZF1 by POM in U937 cells, and this defect could be rescued by wild-type UBE2G1, but not a UBE2G1 enzymatically-dead mutant (C90S, Figure 1C). Knockout of UBE2D3, on the other hand, showed very little effect on the degradation of endogenous IKZF1 (Figure 2C). Thus, we reasoned that there are additional E2 enzyme(s) modulating the degradation of IKZF1 cooperatively with UBE2G1. To identify such E2(s), we evaluated the effect of double knockout of UBE2G1 and one of the 41 E2 enzymes on IKZF1 degradation using a dual gRNA-directed gene knockout approach (Figures 2A and S3). Notably, double knockout of UBE2G1 and UBE2D3 produced more inhibition of POM-induced degradation of ePL-tagged and endogenous IKZF1 than either single knockout alone (Figures 2B and 2C). Combinatorial ablation of UBE2G1 with UBE2E1 or UBE2M also demonstrated subtle but noticeable further inhibition on IKZF1 degradation (Figures S3E and S3I). Although UBE2D2 knockout slightly attenuated the POM-induced ePL-IKZF1 degradation (Figure S2D), double knockout of UBE2D2 and UBE2G1 did not significantly augment the inhibition of ePL-IKZF1 degradation imposed by UBE2G1 single knockout (Figure S3D).

**Figure 2.**
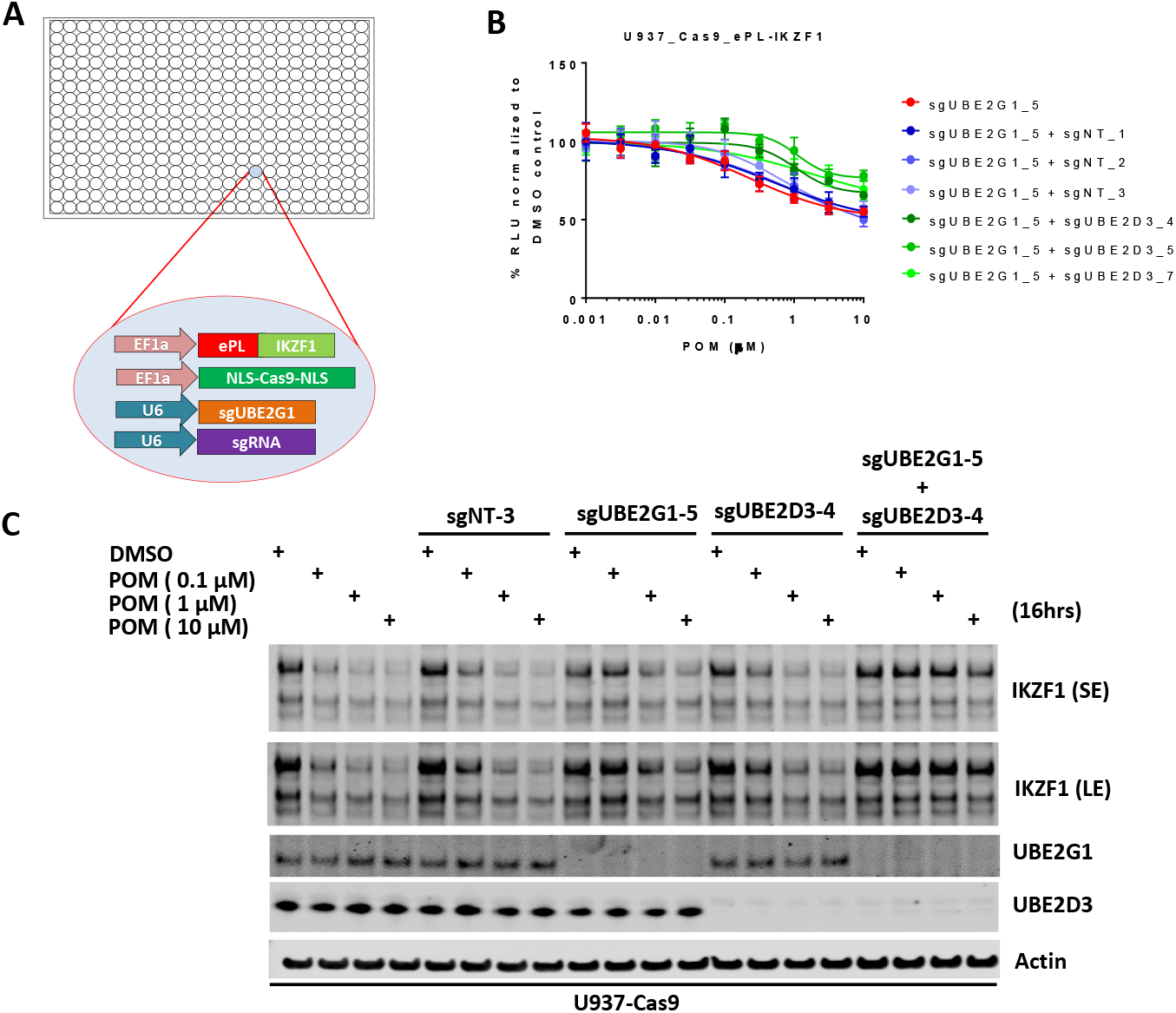
UBE2G1 and UBE2D3 redundantly regulate the pomalidomide-induced degradation of IKZF1. (A) Schematic showing the design of dual-gRNA directed CRISPR screen of E2s regulating the CM-induced degradation of ePL-tagged IKZF1 in 384-well array format. (B) Chemiluminescent measurement of ePL-IKZF1 protein expression level in U937-Cas9_ePL-IKZF1 parental cells or cells expressing UBE2G1-specfic sgRNA alone or in combination with non-targeting or UBE2D3-specific sgRNA. Cells were treated with POM at the indicated concentrations for 16 hours. Data are presented as mean ± SD (n = 4). (C) Immunoblot analysis of U937-Cas9 parental cells or cells expressing non-targeting sgRNA, UBE2G1-specific sgRNA, UBE2D3-specfic sgRNA, or both UBE2G1 and UBE2D3 sgRNAs. Cells were treated with POM at the indicated concentrations for 16 hours. SE, short exposure; LE, long exposure.

To assess the substrate selectivity of UBE2G1, we determined the effect of UBE2G1 knockout on the degradation of IKZF1, its paralog IKZF3 and other well-characterized cereblon neomorphic substrates, triggered by their respective cereblon modulating agents including cereblon-based PROTACs. Ablation of UBE2G1 significantly diminished the degradation of IKFZ1, IKZF3 and ZFP91 by LEN, POM, and CC-220, as well as CK1α degradation by LEN in OPM2, DF15 and MM1S myeloma cells (Figures 3A, S4A, S4B, 7B, S8A and S8B). UBE2G1 loss also reduced the degradation of GSPT1 induced by CC-885 in myeloma cell lines OPM2, DF15 and MM1S (Figures 3B, S4C and S3D), AML cell lines OCI-AML2, U937, MOLM-13 and MV4-11 (Figures S4E-H), as well as 293T human embryonic kidney cells (Figure S4I). The GSPT1 degradation defect conferred by UBE2G1 depletion could also be rescued by UBE2G1 wild-type but not C90S mutant in 293T cells (Figure S4I). UBE2G1 loss also blocked the degradation of Brd4 induced by the cereblon-based BET PROTAC dBET1 (Winter et al., 2015), but not the VHL-based BET PROTAC MZ1 (Zengerle et al., 2015) in 293T cells. UBE2G1 depletion prolonged the protein half-lives of IKZF1 and IKZF3 in OPM2 cells treated with POM. Lastly, UBE2G1 loss did not affect stability of SCF substrates p27 and c-Myc, suggesting that UBE2G1 might be specific to CRL4 (Figure 3C).

**Figure 3.**
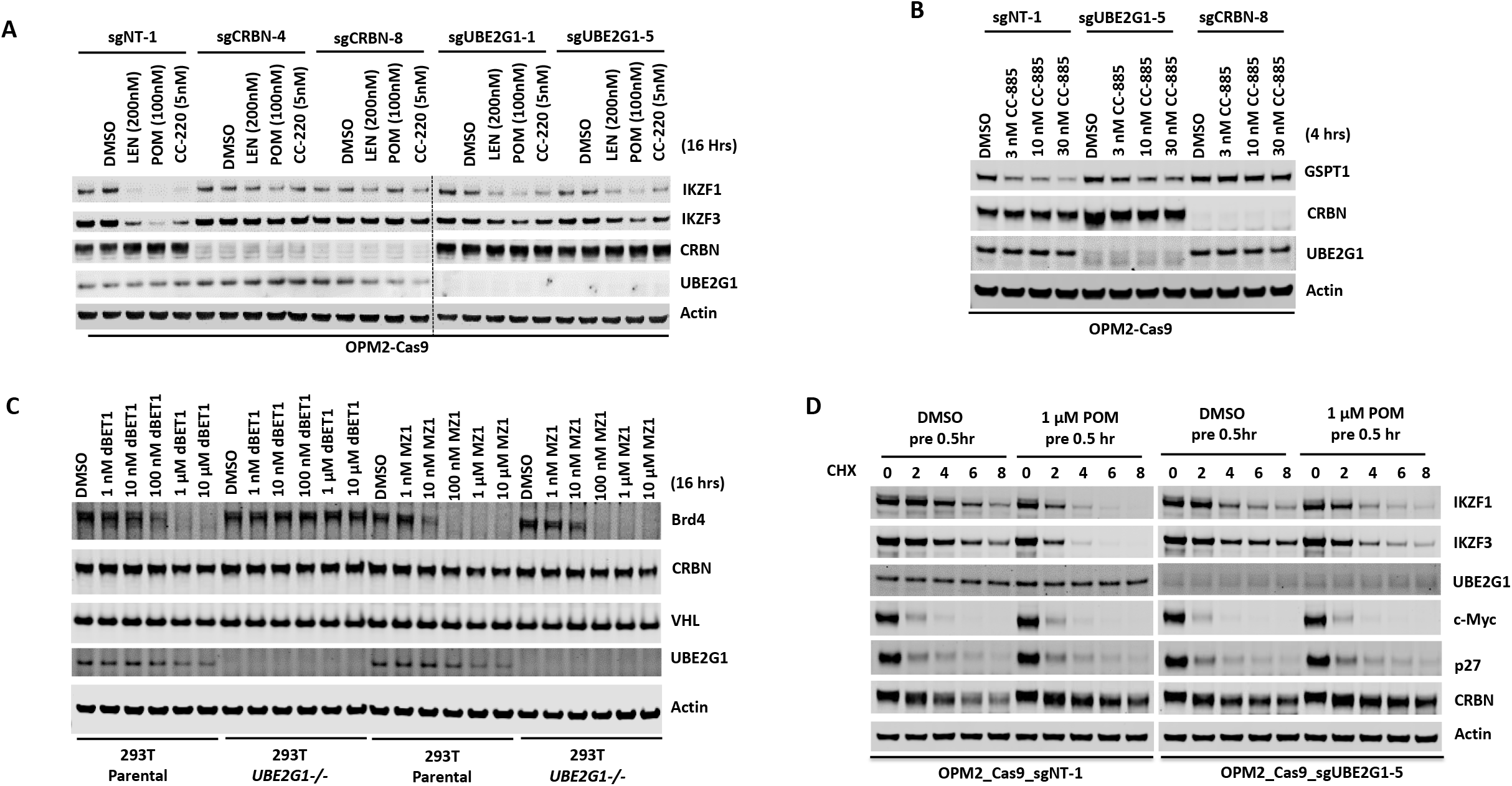
Loss of UBE2G1 blocked the degradation of cereblon neomorphic substrates induced by cereblon-modulating agents. (A and B) Immunoblot analysis of OPM2-Cas9 cells expressing non-targeting, UBE2G1-specific or CRBN-specific sgRNA. Cells were treated with LEN, POM or CC-220 for 16 hrs (A) or CC-885 for 4 hours (B) at the indicated concentrations. (C) Immunoblot analysis of 293T parental or UBE2G1-/- cells treated with Brd4 PROTACs dBET1 or MZ1 at the indicated concentrations for 16 hours. (D) Immunoblot analysis of OPM2 parental or UBE2G1-/- cells treated with 100 μg/ml cycloheximide with or without 1 μM POM pretreatment for half an hour. Cells were harvested at the indicated time points.

### UBE2G1 mediates the ubiquitination of cereblon neomorphic substrates

UBE2G1 and its paralog UBE2G2 share similar domain structures with CDC34. A common structural feature of these three E2 enzymes is an acidic loop (Figure S5A, highlighted with red) in the vicinity of their respective active site cysteines (Figure S5A, highlighted with blue), which facilitates the direct binding with ubiquitin and enables K48-linked ubiquitin chain assembly in the absence of associated E3 ligases (Choi et al., 2015). In association with gp78, an ER membrane bound RING finger E3 ubiquitin ligase, UBE2G2 directly tags misfolded proteins with K48-linked ubiquitin chains preassembled on the catalytic cysteine of UBE2G2, resulting in ER associated protein degradation (ERAD) (Li et al., 2007). Although it has been shown that UBE2G1 and UBE2G2 redundantly mediate the destabilization of the CRL4^cdt2^ substrate Cdt1 in response to UV irradiation, the direct transfer of ubiquitin to Cdt1 by UBE2G1 or UBE2G2 has not been demonstrated (Shibata et al., 2011).

Next, we employed a reconstituted *in vitro* ubiquitination assay to address the role of UBE2G1 and UBE2D3 in ubiquitination of IKZF1 and GSPT1 induced by POM and CC-885, respectively. We monitored the production of ubiquitin conjugates of IKZF1 and GSPT1 catalyzed by UBE2D3 alone, UBE2G1 alone, or in combination, in the presence of Ube1 (E1), Cul4-Rbx1, cereblon-DDB1, ubiquitin and ATP with or without POM or CC-885. UBE2D3 alone produced ubiquitin conjugates of IKZF1 or GSPT1 mainly with a single or di-ubiquitin moiety in a POM-or CC-885-dependent manner (Figures 4A and 4B). By contrast, UBE2G1 alone failed to display any ubiquitin conjugating activity for both IKZF1 and GSPT1 (Figures 4A and 4B, lanes 3 and 4). However, when combined with UBE2D3, UBE2G1 significantly promoted the extent of ubiquitination of both substrates (Figures 4A and 4B, lanes 2 and 6). Moreover, the ubiquitin conjugates of IKZF1 or GSPT1 formed with both UBE2G1 and UBE2D3, but not UBE2D3 alone, were exclusively K48-linked, because the wild-type ubiquitin used in the reconstituted ubiquitination reaction could be functionally replaced by ubiquitin mutant K48-only (with 6 remaining lysine residues mutated to arginine), but not by K48R (remaining lysine residues were intact), and the ubiquitination pattern of IKZF1 or GSPT1 catalyzed by UBE2D3 exhibited no obvious difference between wild-type ubiquitin and K48-only or K48R mutant (Figures 4A and 4B, lane 2; Figures 4C and 4D, lanes 2 and 8).

**Figure 4.**
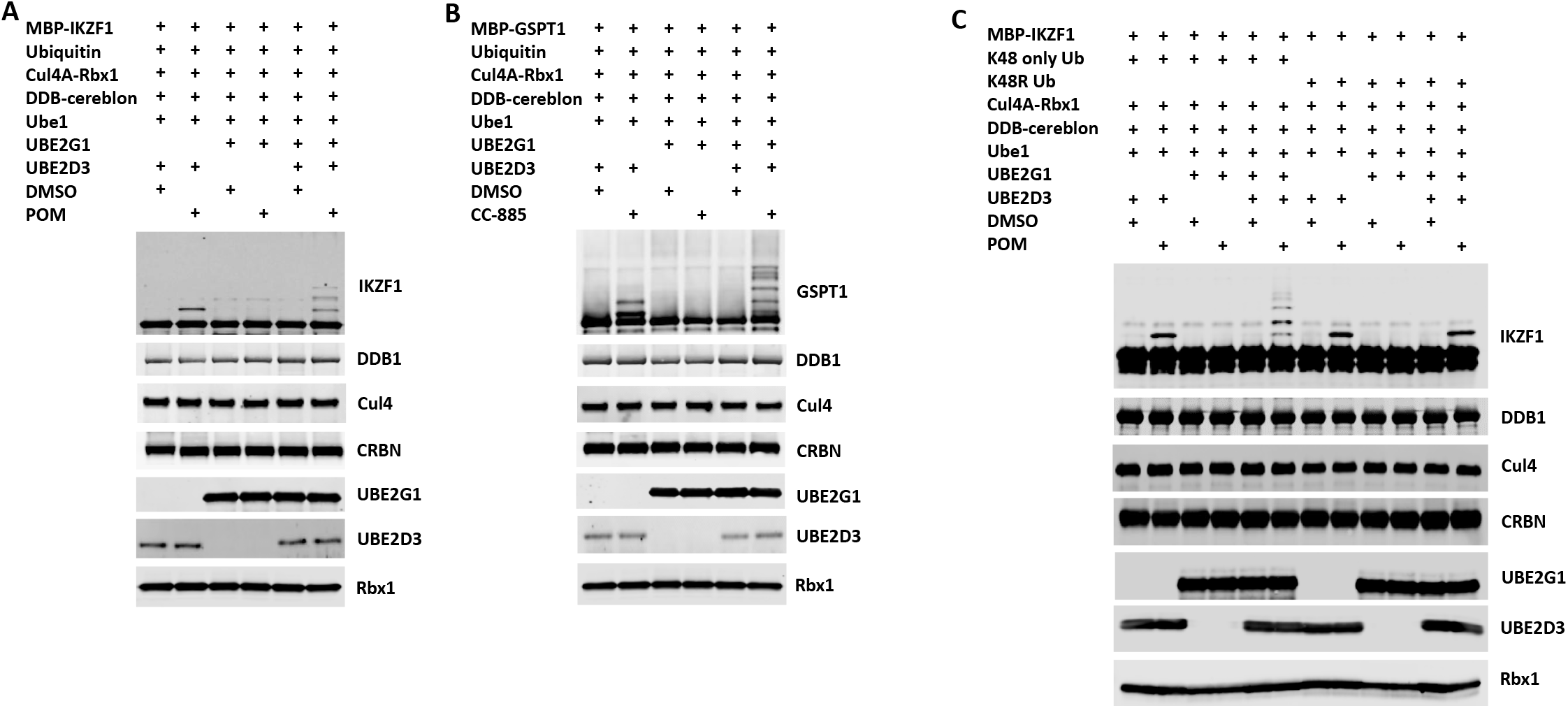
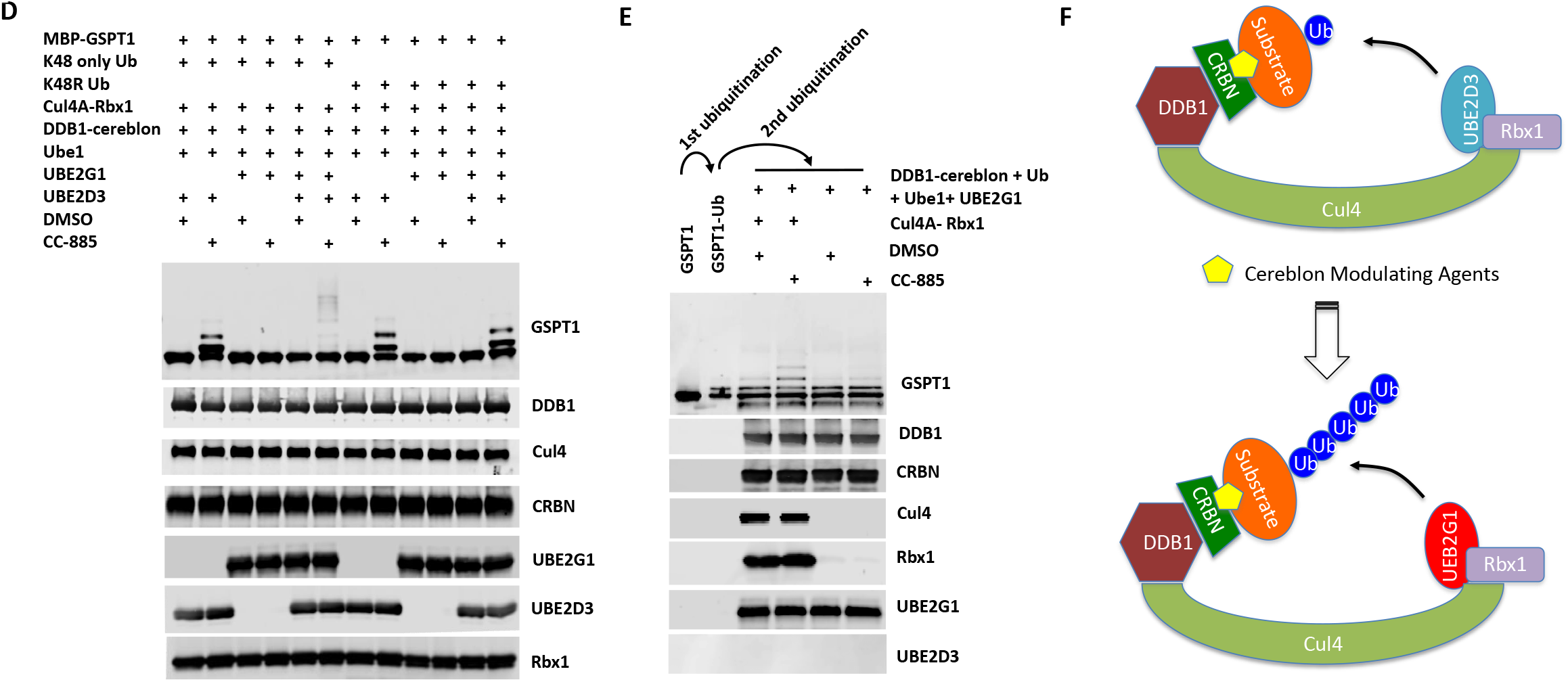
UBE2G1 and UBE2D3 sequentially catalyze the *in vitro* ubiquitination of IKZF1 and GSPT1 in the presence of pomalidomide and CC-885, respectively. (A-D) In vitro ubiquitination of IKZF1 (A and C) and GSPT1 (B and D) MBP fusion proteins by recombinant CRL4CRBN complex. Recombinant protein products as indicated were incubated with or without 80 μM POM (A and C) or 80 μM CC-885 (B and D) in the ubiquitination assay buffer containing 80 mM ATP at 30°C for 2 hours, and then analyzed by immunoblotting. (E) Sequential in vitro ubiquitination of GSPT1 by recombinant CRL4CRBN complex. MBP-GSPT1 recombination protein was incubated with Ube1, UBE2D3, Cul4-Rbx1, DDB1-cereblon, Ubiquitin, ATP and CC-885 in the ubiquitination assay at 30 °C for 4 hours. After purification over size-exclusion chromatography, pre-ubiquitinated MBP-GSPT1 protein was then incubated with Ube1, DDB1-cereblon, Ubiquitin, ATP and UBE2G1 with or without CC-885 or Cul4A-Rbx1 in the ubiquitination assay at 30 °C for 2 hours, followed by immunoblot analysis. (F) Schematic showing the sequential ubiquitination of CRBN neomorphic substrates by UBE2D3 and UBE2G1.

To further explore the mechanism underlying the cooperativity between UBE2G1 and UBE2D3, we separated the ubiquitination reaction of GSPT1 into two steps. First, following GSPT1 ubiquitination by UBE2D3 alone, we isolated GSPT1 ubiquitin conjugates from the rest of reaction components using a size-exclusion column (Figure 4E, lanes 1 and 2). We then incubated the purified GSPT1 ubiquitin conjugates with UBE2G1, Ube1 (E1), cereblon-DDB1, ubiquitin and ATP with or without CC-885 and Cul4-Rbx1. We found that UBE2G1 was capable of catalyzing the further ubiquitination of GSPT1 only with prior-conjugated ubiquitin (Figure 4E, lanes 1-4, note the conversion of the mono-ubiquitinated GSPT1 into di- or tri-ubiquitinated forms), and this action required the presence of CC-885 and Cul4A-Rbx1 (Figures 4E).

To rule out the possibility that the UBE2G1 function observed above is simply an artifact of bacterial recombinant protein, we reconstituted the ubiquitination reaction using FLAG-tagged UBE2G1 and FLAG-tagged UBE2D3 proteins purified from 293T *UBE2G1-/-* cells, in which ectopic overexpression of FLAG-tagged UBE2G1 and UBE2D3, but not their respective enzymatically-dead mutant, could partially rescue the defect in CC-885-induced GSPT1 degradation elicited by UBE2G1 loss (Figure S5B). In agreement with our previous findings, GSPT1 ubiquitination catalyzed *in vitro* by FLAG-UBE2G1 and FLAG-UBE2D3, alone or in combination, was similar to what was observed with bacterial recombinant UBE2G1 and UBE2D3 (Figure S5C). In addition, we examined the *in vivo* function of UBE2G1 and UBE2D3 in the regulation of POM-induced ubiquitination of IKZF1 ectopically expressed in 293T cells. Ablation of both UBE2G1 and UBE2D3 significantly reduced, but did not completely block, the mono- and polyubiquitination of IKZF1 induced by POM, indicating the existence of additional E2 enzymes with redundant function (Figure 5A and S6A). Reintroduction of UBE2G1 alone via transient transfection in 293T *UBE2G1-/- ; UBE2D3-/-* cells dramatically enhanced the extent of POM-induced IKZF1 polyubiquitination (Figure 5B, lanes 5-8 vs 1-4). Transient overexpression of UBE2D3 alone promoted IKZF1 monoubiquitination as well as polyubiquitination but to a lesser extent (Figure 5B, lanes 9-12 vs 1-4). Overexpression of both achieved an additive or possibly synergistic, effect on IKZF1 ubiquitination (Figure 5B, lanes 13-16 vs 5-12, note the conversion of monoubiquinated IKZF1 into its polyubiquitinated forms). In keeping with the *in vitro* finding (Figure S5C), IKZF1 was mainly mono-ubiquitinated in 293T *UBE2G1-/-* cells following POM treatment, and UBE2G1 wild-type but not C90S mutant could restore IKZF1 polyubiquitination (Figure 5C and S6C).

**Figure 5.**
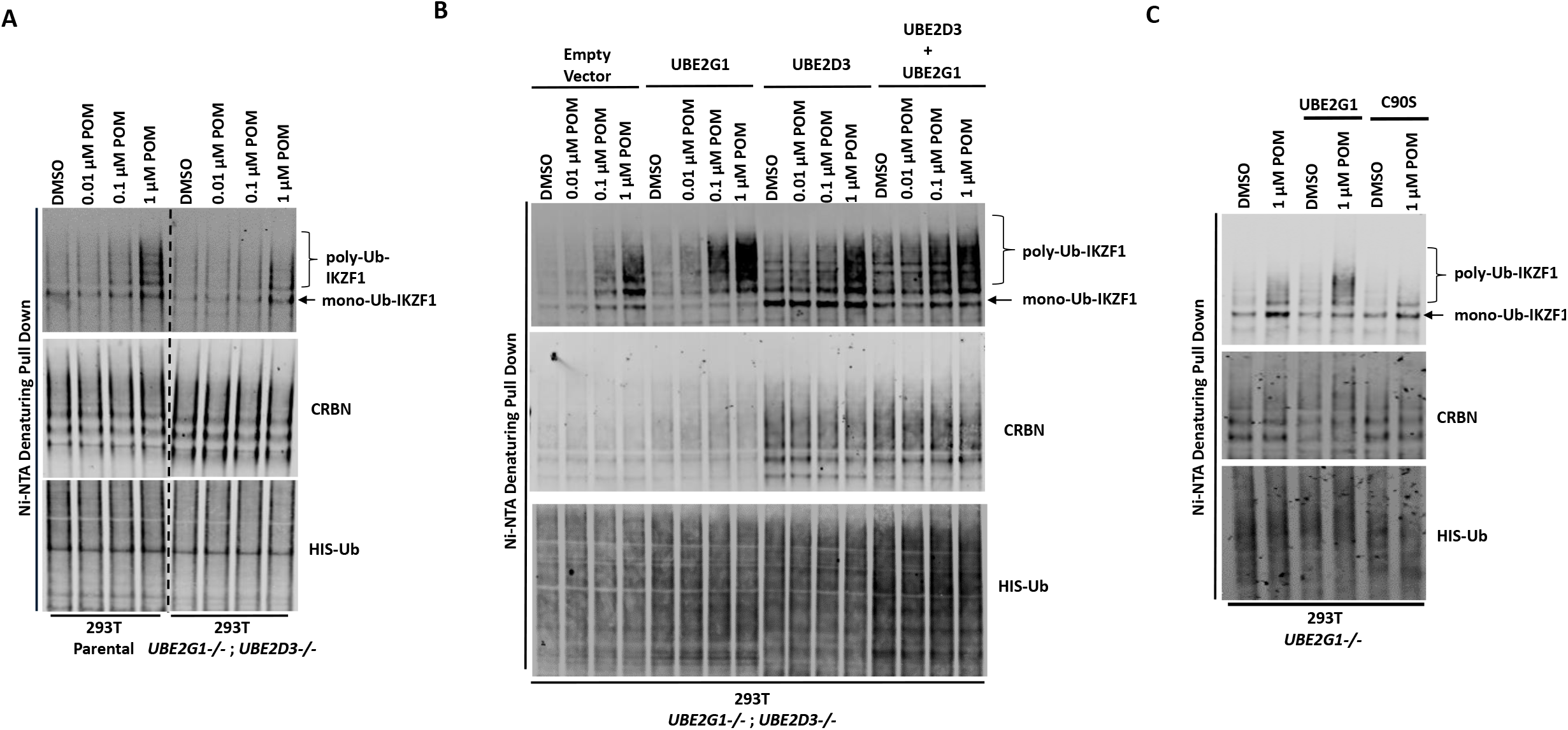
UBE2G1 and UBE2D3 cooperatively promote the *in vivo* ubiquitination of IKZF1. (A and B) 293T parental and UBE2G1-/-;UBE2D3-/- (clone 4) cells were transiently transfected with plasmids expressing cereblon, V5-tagged IKZF1 and 8xHis-Ub with or without UBE2G1, UBE2D3 or both. (C) 293T parental and UBE2G1-/- (clone 13) cells were transiently transfect with plasmids expressing cereblon, IKZF1-V5, 8xHis-Ub with or without UBE2G1 wild-type or C90S mutant. In (A), (B) and (C), 48 hours after transfection, cells were treated with MG-132 (10 μM) and POM at the indicated concentrations for additional 8 hours. Ubiquitinated protein products enriched with magnetic nickel sepharose were subjected to immunoblot analysis. Immunoblot analysis of whole cell extracts showing equal input proteins is shown in Supplemental Figures 6A, 6B and 6C.

### UBE2G1 loss confers resistance to cereblon modulating agents

Since UBE2G1 depletion significantly attenuated the degradation of all cereblon neomorphic substrates, we reasoned that UBE2G1 protein downregulation, gene deletion or mutation might lead to reduced CRL4^CRBN^ activity, thereby leading to resistance to cereblon modulating agents. To test this hypothesis, we surveyed the protein expression level of UBE2G1 in myeloma cell lines with variable sensitivity to LEN and POM (Figure 6A). LEN sensitive cell lines MM1S, OPM2, DF15 and NCI-H929 displayed higher expression level of UBE2G1 than LEN medium-sensitive cell line ANABL-6 and LEN resistant cell lines EJM, L363, SKMM2, CAG and ARH-77, and more strikingly UBE2G1 expression in SKMM2 was undetectable (Figure 6B). As expected, reintroduction of UBE2G1 wild-type but not C90S mutant significantly augmented the antiproliferative effect of both LEN and POM, which was linked to the enhanced degradation of IKZF1 and IKZF3 (Figures 6C and 6D). UBE2G1 loss also conferred resistance to CC-885 in OCI-AML2, U937, MOLM-13, and MV4-11 AML cells (Figures S7A-D), as well as 293T cells, and this defect could be rescued by UBE2G1 wild-type but not C90S mutant (Figure S7E). Moreover, UBE2G1 deletion conferred resistance to BET PROTAC dBET1 but not MZ1 in 293T cells. Lastly, UBE2G1 knockout in DF15, MM1S and OPM2 myeloma cells conferred significant resistance to LEN and POM. Importantly, these cells were only partially resistant to CC-220, in keeping with the increased efficiency in triggering IKZF1 and IKZF3 degradation (Figures 7A, 7B and S8A-D). These results indicate that UBE2G1 deficiency may be a key differentiator in the clinical success of cereblon modulating agents.

**Figure 6.**
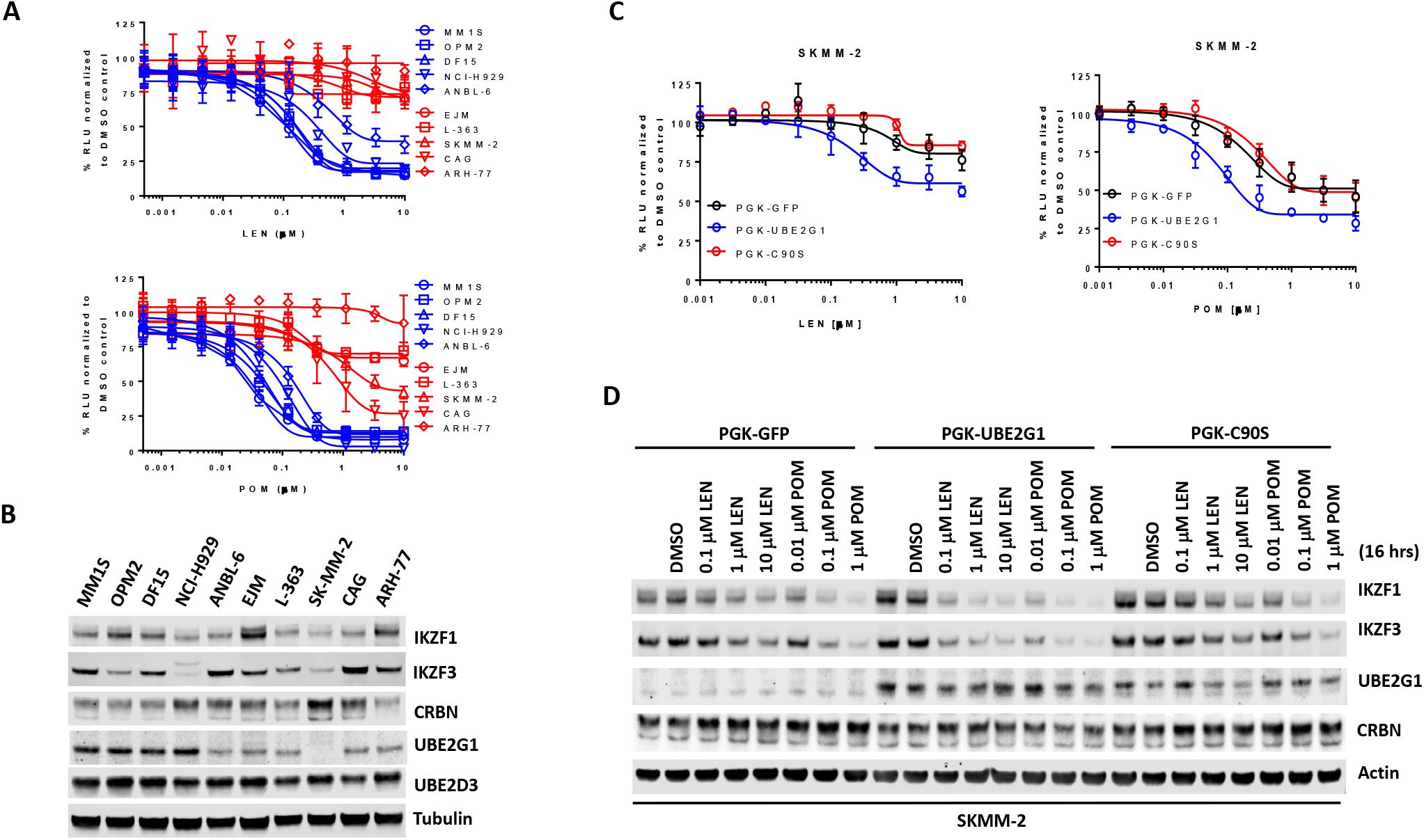
UBE2G1 loss confers resistance to lenalidomide and pomalidomide in myeloma cell lines. (A) Effect of lenalidomide (top panel) and pomalidomide (bottom panel) on proliferation of myeloma cell lines. Cell proliferation was determined by CTG. Data are presented as mean ± SD (n=3). (B) Immunoblot analysis of myeloma cell lines. (C and D) Proliferation (C) and immunoblot analysis (D) of SKMM2 cells transduced with lentiviral vectors encoding GFP, UBE2G1 and UBE2G1-C90S. Cells were treated with DMSO vehicle control, LEN or POM at the indicated concentrations for 5 days (C) or 16 hours (D). In (C), cell proliferation was determined by CTG, and data are presented as mean ± SD (n=3).

**Figure 7.**
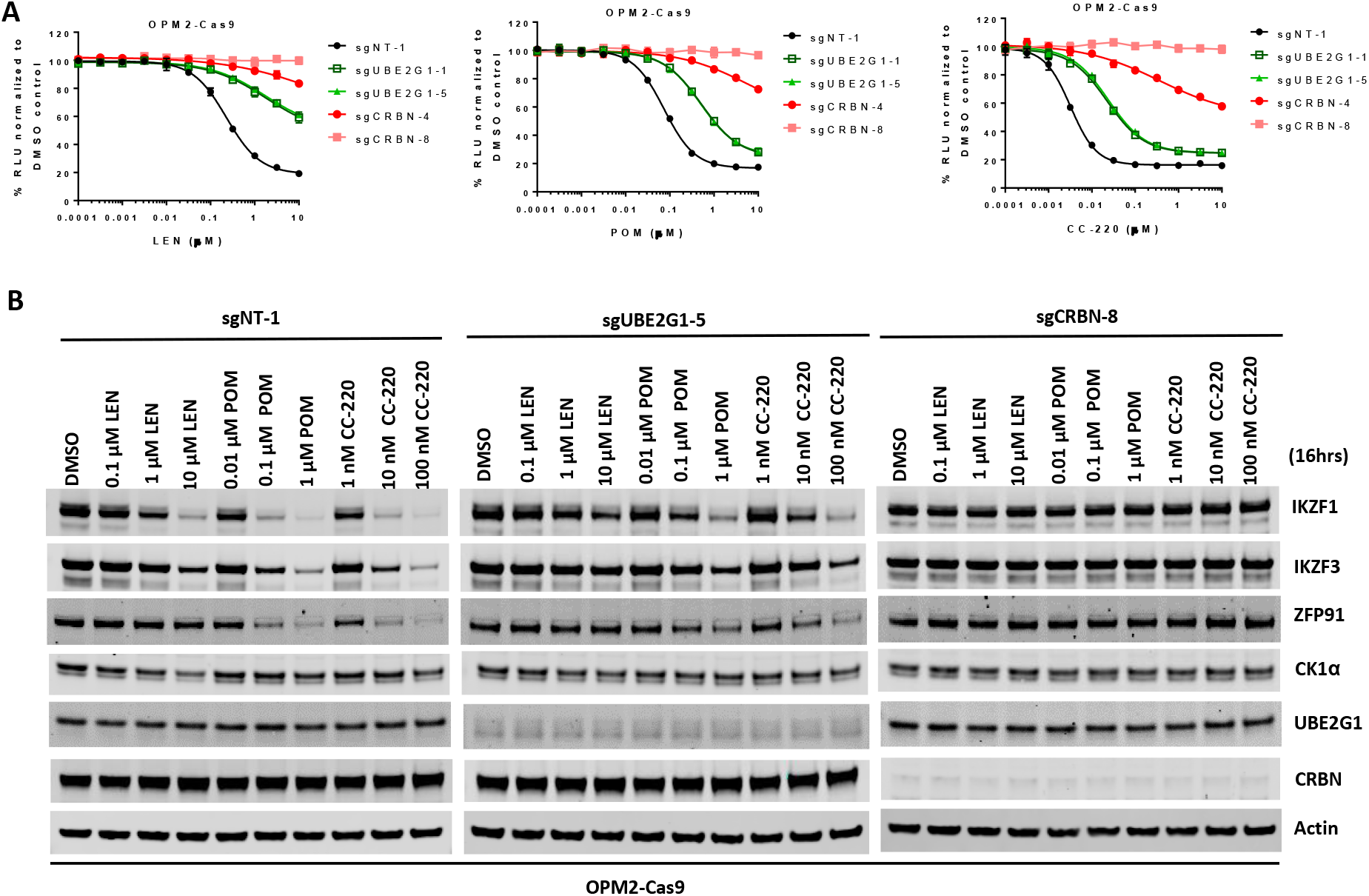
UBE2G1-deficient OPM2 myeloma cells are resistant to lenalidomide and pomalidomide but remain sensitive to CC-220 at clinical relevant concentrations. (A and B) Cell proliferation (A) and immunoblot analysis (B) of OPM2-Cas9 cells transduced with lentiviral vectors expressing non-targeting, UBE2G1-specific or CRBN-specific sgRNAs. Cells were treated DMSO vehicle control, LEN, POM or CC-220 at the indicated concentrations for 5 days (A) or 16 hours (B). In (A), cell proliferation was determined by CTG, and data are presented as mean ± SD (n=3).

## Discussion

The sequential recruitment of two functionally distinct E2s by a single E3 is a general mechanism for substrate ubiquitination conserved from yeast to human (Rodrigo-Brenni and Morgan, 2007) (Wu et al., 2010) (Kleiger and Deshaies, 2016). In this work, we provide evidence that CRL4^CRBN^ deploys the same mechanism to mark its neomorphic substrates with K48-linked poly-ubiquitin chains for proteasomal degradation. Mechanistically, UBE2D3 transfers the first ubiquitin onto the lysine residue(s) of the cereblon neomophic substrate, thereby enabling UBE2G1 to assemble the K48-linked ubiquitin chains onto the initial anchor ubiquitin (Figure 4F). This orchestrated action between UBE2D3 and UBE2G1 closely resembles the cooperativity of UBE2D3 and Cdc34 in promoting IκBα polyubiquitination mediated by SCF^βTRCP2^ (Wu et al., 2010), except that unlike Cdc34, UBE2G1 does not possess the ability to transfer ubiquitin onto substrates without prior ubiquitin conjugation. UBE2G1 was known to produce K48-linked poly-ubiquitin chains in the absence of an E3 ubiquitin ligase (Choi et al., 2015), but UBE2G1 cannot promote GSPT1 ubiquitination in the absence of CC-885 or Cul4A-Rbx1 (Figure 4E), indicating that close proximity of UBE2G1 to cereblon neomorphic substrates bridged by CRL4^CRBN^ is required to increase the processivity of UBE2G1 at physiological concentrations. This provides an explanation to how cereblon modulating agents can induce effective substrate degradation via K48-linked polyubiquitination, as well as speaks to the potential role of UBE2G1 in mediating the ubiquitination and degradation of other CRL4 cognate and/or neomophic substrates. This is supported by the impaired degradation of p21 and RBM39 induced by UV irradiation and E7070 treatment, respectively, in 293T *UBE2G1-/-* cells (Figures S9A and S9B).

Although UBE2D3 and UBE2G1 cooperatively regulate the ubiquitination of cereblon neomorphic substrates, ablation of UBE2D3 exhibited very little impact on substrate degradation as compared to loss of UBE2G1, suggesting that additional E2s might fulfill the role of UBE2D3, e.g., other UBE2D family proteins with a high degree of sequence homology including UBE2D1/UbcH5a, UBE2D3, and UBE2D4 (Figure S10A). This idea is supported by the following observations. First, just like UBE2D3, UBE2D1 and UBE2D2 acted synergistically with UBE2G1 in catalyzing the ubiquitination of GSPT1 in the presence of CC-885, whereas cooperativity among UBE2D1, UBE2D2 and UBE2D3 cannot be detected (Figure S10B). Second, UBE2D1 and to a lesser extent UBE2D2 were far more efficient than UBE2D3 in catalyzing CC-885-dependent GSPT1 polyubiquitination with or without the help of UBE2G1 (Figure S10B, lanes 2, 4 and 6). Third, UBE2D2 knockout impaired the POM-induced degradation of ePL-tagged IKZF1 (Figure S2D). Although the synergistic interaction between UBE2D2 and UBE2G1 was not evident in the dual-gRNA CRISPR screen (Figure S3D), their cooperativity might still exist but fall below the detection limit of this assay. Regardless, it is more likely that UBE2D2 could generate extensive substrate polyubiquitination in the absence of UBE2G1 *in vivo*, leading to substrate degradation. Taken together, we speculate that the CM-induced proteolysis of cereblon neomorphic substrates is both redundantly and cooperatively regulated by UBE2D family proteins and UBE2G1, with UBE2G1 playing a key role in enhancing the rate and extent of K48-linked substrate ubiquitination, resulting in rapid and efficient substrate degradation.

Although loss of UBE2G1 conferred resistance to LEN and POM in human myeloma cell lines, it remains to be seen whether UBE2G1 deficiency occurs in human myeloma patients with inherent or acquired resistance to IMiD drug treatment, especially those with normal cereblon expression. The myeloma cell line SKMM2, which lost both copies of *UBE2G1* gene (based on CCLE gene copy number characterization), and has undetectable UBE2G1 protein expression (Figure 6B), was derived from a human myeloma patient who never received any prior treatment with IMiD drugs (Eton et al., 1989), warranting the further clinical evaluation of UBE2G1 activity in myeloma patients. Given that UBE2G1 inactivation conferred resistance to all CMs tested and also to cereblon-based PROTACs, patient stratification approaches based on UBE2G1 status might be applicable to the development of IMiD drugs and other novel cereblon modulating agents for a variety of human diseases. Lastly, CC-220, a novel CM that targets IKZF1 and IKZF3 for degradation much more effectively than does LEN or POM, retained strong antitumor activity at clinically achievable concentrations (Schafer et al., 2018) in UBE2G1-deficient myeloma cells, suggesting that human patients with resistance to CM drugs owing to diminished UBE2G1 function may be responsive to next-generation CMs that possess higher efficiency and/or potency for degrading the same target protein.

## Acknowledges

The authors thank members of the early drug discovery team at Celgene for helpful discussion and technical support.

## Material and Methods

### Purification of *In Vitro* Ubiquitination Assay Components

#### Cereblon-DDB1 purification

ZZ-domain-6xHis-thrombin-tagged human cereblon (amino acids 40 – 442) and full length human DDB1 were co-expressed in SF9 insect cells in ESF921 medium (Expression Systems), in the presence of 50 uM zinc acetate. Cells were resuspended in buffer containing 50 mM Tris-HCl pH 7.5, 500 mM NaCl, 10 mM imidazole, 10% glycerol, 2 mM TCEP, 1X Protease Inhibitor Cocktail (San Diego Bioscience), and 40,000 U Benzonase (Novagen), and sonicated for 30 s. Lysate was clarified by high speed centrifugation at 30,000 rpm for 30 minutes, and clarified lysate was incubated with Ni-NTA affinity resin (Qiagen) for 1 hour. Complex was eluted with buffer containing 500 mM imidazole, and the ZZ-domain-6xHis tag removed by thrombin cleavage (Enzyme Research) overnight, combined with dialysis in 10 mM imidazole buffer. Cleaved eluate was incubated with Ni-NTA affinity resin (Qiagen), and the flow-through diluted to 200 mM NaCl for further purification over an ANX HiTrap ion exchange column (GE Healthcare). The ANX column was washed with 10 column volumes 50 mM Tris-HCl pH 7.5, 200 mM NaCl, 3 mM TCEP, followed by 10 column volumes of 50 mM Bis-Tris pH 6.0, 200 mM NaCl, 3 mM TCEP, and the cereblon-DDB1 peak eluted at 210 mM NaCl. This peak was collected and further purified by size exclusion chromatography using a Sephacryl S-400 16/60 column (GE Healthcare) in buffer containing 10 mM HEPES pH 7.0, 240 mM NaCl, and 3 mM TECP. The cereblon-DDB1 complex was concentrated to 30 mg/mL.

#### Cul4-Rbx1 purification

Human full length Cul4A and Rbx1 were co-expressed in SF9 insect cells. Cells were resuspended in buffer containing 50 mM Tris-HCl pH 7.5, 500 mM NaCl, 10 mM imidazole, 10% glycerol, 2 mM TCEP, 1X Protease Inhibitor Cocktail (San Diego Bioscience), and 40,000 U Benzonase (Novagen), and sonicated for 30 s. Lysate was clarified by high speed centrifugation at 30,000 rpm for 30 minutes, and clarified lysate was incubated with Ni-NTA affinity resin (Qiagen) for 1 hour. Complex was eluted with buffer containing 500 mM imidazole, and concentrated for size exclusion chromatography. Complex was further purified over an S200 Superdex 200 16/600 column (GE Healthcare) in buffer containing 200 mM NaCl, 50 mM Tris pH 7.5, 3 mM TCEP, and 10% glycerol.

#### Substrate purification

MBP-IKZF1 (amino acids 140-168) or MBP-GSPT1 (amino acids 437-633) was expressed in *E.coli* BL21 (DE3) Star cells (Life Technologies) using 2XYT media (Teknova). Cells were induced at OD600 0.6 for 18 hours at 16 °C, with 150 uM zinc acetate added upon induction for IKZF1 expression. Cells were pelleted and resuspended in buffer containing 200 mM NaCl, 50 mM Tris pH 7.5, 3 mM TCEP, 10% glycerol, 150 μM zinc acetate, 0.01 mg/mL lysozyme (Sigma), 40,000 U benzonase (Novagen), and 1X protease inhibitor cocktail (San Diego Bioscience). Resuspended cells were frozen, thawed for purification, and sonicated for 30 s before high speed centrifugation at 30,000 rpm for 30 minutes. Clarified lysate was incubated with maltose affinity resin (NEB) at 4 °C for 1 hour before beads were washed. Protein was eluted with buffer containing 200 mM NaCl, 50 mM Tris pH 7.5, 3 mM TCEP, 10% glycerol, 150 μM zinc acetate, and 10 mM maltose. Eluate was concentrated and further purified by size exclusion chromatography over a Superdex 200 16/600 column (GE Healthcare) in buffer containing 200 mM NaCl, 50 mM Tris pH 7.5, 3 mM TCEP, 10% glycerol, and 150 μM zinc acetate.

#### UBE2G1 purification from *E. coli*

Human full length UBE2G1 with an N-terminal 6XHis-thrombin tag was expressed in *E.coli* BL21 (DE3) Star cells (Life Technologies) using 2XYT media (Teknova). Cells were induced at OD_600_ 0.6 for 18 hours at 16 °C. Cells were pelleted and resuspended in buffer containing 50 mM Tris pH7.5, 250 mM NaCl, 3mM TCEP, 1X Protease Inhibitor Cocktail (San Diego Bioscience), 20 mM imidazole, and 40,000 U Benzonase (Novagen), and sonicated for 3 times for 30 s. Lysate was clarified by high speed centrifugation at 30,000 rpm for 30 minutes, and clarified lysate was incubated with Ni-NTA affinity resin (Qiagen) for 1 hour at 4C. The protein was eluted with buffer containing 500 mM imidazole. The 6XHis tag was then removed by thrombin cleavage (Enzyme Research) overnight, combined with dialysis into 50 mM Tris pH7.5, 250 mM NaCl, 3mM TCEP, 1X Protease Inhibitor Cocktail (San Diego Bioscience), 20 mM imidazole. The cleaved protein was then loaded onto a 5ml HiTrap Ni column (GE Healthcare), and cleaved protein collected in the flow-through. The flow-through was then concentrated and further purified by size exclusion chromatography over a Superdex 75 16/600 column (GE Healthcare) in buffer containing 20mM Tris pH7.5, 150mM NaCl, 1mM DTT, and concentrated to 25 μM.

#### E2 purification from 293T cells

FLAG tagged UBE2D3 wild-type and C85S mutant, and FLAG-tagged UBE2G1 wild-type and C905S mutant were purified from 293T *UBE2G1-/-* cell lines stably expressing the respective protein. A pellet of ~5000 million cells expressing a FLAG-tagged E2 was re-suspended in buffer containing 50 mM Tris-HCl pH 7.5, 250 mM NaCl, 1 mM TCEP, 1X Protease Inhibitor Cocktail (San Diego Bioscience), and Phosphatase inhibitor cocktail (Sigma (Roche), REF 04 906 837 001), and sonicated for 15s. Lysate was clarified by high speed centrifugation at 30,000 rpm for 30 minutes, and clarified lysate was incubated with anti-FLAG M2 Affinity Gel (A2220 Sigma) for 2 hour. The E2 was eluted with buffer containing 50 mM Tris pH7.5, 250 mM NaCl, 0.15 mg/ml FLAG peptide (3X FLAG Peptide F4799, Sigma Aldrich). The protein was then dialyzed into 4 L of buffer containing 50 mM Tris pH 7.5 and 250 mM NaCl for one hour twice. The protein was then concentrated and further purified by size exclusion chromatography over a Superdex 75 16/600 column (GE Healthcare) in buffer containing 20mM Tris pH 7.5, 150 mM NaCl. All 4 proteins were then concentrated to 25 μM final concentration.

### *In Vitro* Ubiquitination Assays

Purified E1, E2, ubiquitin, Cul4A-Rbx1, and cereblon-DDB1 proteins were used to reconstitute the ubiquitination of MBP-fused GSPT1 or IKZF1 substrates *in vitro.* Purified recombinant human Ube1 E1 (E-305), UbcH5a/UBE2D1 (E2-616-100), UbcH5b/UBE2D2 (E2-622-100), UbcH5c/UBE2D3 (E2-627-100), wild-type ubiquitin (U-100H), K48R ubiquitin (UM-K48R-01M), and K48-only ubiquitin (UM-K480-01M) were purchased from R&D systems. For the ubiquitination of IKZF1 or GSPT1 shown in Figures 4A, 4B S5C and S10B, reaction components were mixed to final concentrations of 80 mM ATP, 1.5 μM Ube1, 275 μM Ub, 2 μM Cul4-Rbx1, 2 μM cereblon-DDB1, 5 uM IKZF1 (a.a. 140-168) or 5 μM MBP-GSPT1 (a.a. 437-633) as indicated, and then 5 μM UBE2D1, 5 μM UBE2D2, 5 μM UBE2D3 (purified from either from E.coli or human cells), or 7.5 μM UBE2G1 (purified from either from E.coli or human cells), was added alone or in combination, as indicated. Reactions were incubated in the presence of either DMSO or 80 μM compound (pomalidomide or CC-885) in ubiquitination assay buffer (20 mM HEPES pH 7.5, 150 mM NaCl, 10 mM MgCl2). To start the reactions, E1, E2, ATP and ubiquitin were pre-incubated for 30 minutes, and separately MBP-substrate, CRBN-DDB1, Cul4-Rbx1, and compound were pre-incubated for 5 minutes at room temperature, before ubiquitination reactions were started by mixing the two pre-incubations. Reactions were incubated at 30°C for 2 hours before separation by SDS-PAGE followed by immunoblot analysis using anti-MBP antibody (MBP-probe R29.6, Santa Cruz). For the ubiquitin mutant reactions shown in Figures 4C and 4D, 275 μM K48-only or K48R ubiquitin was substituted for wild-type ubiquitin as indicated, and the E. coli-purified UBE2G1 was used. Reactions were incubated at 30°C for 2 hours before separation by SDS-PAGE followed by immunoblot analysis using anti-MBP antibody (MBP-probe R3.2, Santa Cruz).

For the ubiquitination of a pre-ubiquitinated substrate shown in figure 4E, MBP-GSPT1 (a.a. 437-633) was incubated with 80 mM ATP, 3 μM Ube1, 600 μM Ub, 4 μM Cul4-Rbx1, 4 μM cereblon-DDB1, 5 μM UBE2D3 (purified from E. coli), and 80 uM CC-885 for 4 hours before separation of the reaction over a 10/300 S200 GL (GE 17-5175-01) size exclusion chromatography column to separate the substrate from the rest of the ubiquitination reaction components. 1.25 μM purified MBP-GSPT1 was then used as the substate in ubiquitnation reactions including 80 mM ATP, 1.5 μM Ube1, 600 μM Ub, 2 μM Cul4-Rbx1, 2 μM cereblon-DDB1, 80 μM CC-885, and 7 μM UBE2G1 (purified from E. coli). Reactions were incubated at 30°C for 2 hours before separation by SDS-PAGE followed by immunoblot analysis using anti-MBP antibody (MBP-probe R29.6, Santa Cruz).

### Cell Culture and Materials

Human embryonic kidney cell line 293T (Clontech) was maintained in Dulbecco’s Modified Eagle’s medium (DMEM; Invitrogen) supplemented with 10% fetal bovine serum (FBS; Invitrogen), 1x sodium pyruvate (Invitrogen), 1x non-essential amino acids (Invitrogen), 100 U/mL penicillin (Invitrogen), and 100 μg/mL streptomycin (Invitrogen). Acute myeloid leukemia cell lines U937, MOLM-13, and MV4-11 and myeloma cell line MM1S were purchased from American Tissue Culture Collection (ATCC). Acute myeloid leukemia cell line OCI-AML2 cell line and myeloma cell lines OPM2 was purchased from Deutsche Sammlung von Mikroorganismen und Zellkulturen GmbH (DSMZ). Myeloma cell line DF15 was obtained from Dr John Shaughnessy (University of Arkansas, Little Rock, AR, USA). U937, MOLM-13, OPM2, MM1S and DF15 cell lines were maintained in Roswell Park Memorial Institute (RPMI) 1640 tissue culture medium (Invitrogen) supplemented with 10% FBS, 1x sodium pyruvate, 1x non-essential amino acids, 100 U/mL penicillin, and 100 μg/mL streptomycin. MV4-11 cell line was maintained in Iscove’s Modified Dulbecco’s medium (IMDM; (Invitrogen) supplemented with 10% FBS, 1x sodium pyruvate, 1x non-essential amino acids, 100 U/mL penicillin, and 100 μg/mL streptomycin. OCI-AML2 cell line was maintained in minimal essential medium (MEM; Invitrogen) supplemented with 10% FBS, 1x sodium pyruvate, 1x non-essential amino acid, 100 U/mL penicillin, and 100 μg/mL streptomycin. All cell lines were cultured at 37°C with 5% CO2 in the relevant media mentioned above.

### Plasmids

UBE2G1 and UBE2D3 complimentary deoxyribonucleic acid (cDNA) clones were purchased from Dharmacon. The coding regions of UBE2G1 and UBE2D3 were polymerase chain reaction (PCR)-amplified and shuttled into pDONR223 via BP (attB and attP) recombination to generate pDONR223-UBE2G1, pDONR223-FLAG-UBE2G1, and pDONR223-FLAG-UBE2D3. Site-directed mutagenesis using overlapping PCR was then carried out to generate pDONR223-UBE2G1-CR (CRISPR resistant), pDONR223-UBE2G1-C90S-CR, pDONR223-FLAG-UBE2G1-CR, and pDONR223-FLAG-UBE2G1-C90S-CR, and pDONR223-FLAG-UBE2D3-C85S. Next, gateway donor vectors pDONR223-UBE2G1-CR and pDONR223-UBE2G1-C90S-CR were shuttled into plenti-Ubcp-gateway-IRES-Pur or plenti-PGK-gateway-IRES-Pur via LR (attL and attR) recombination to generate plenti-Ubcp-UBE2G1-CR-IRES-Pur, plenti-Ubcp-UBE2G1-C90S-CR-IRES-Pur, plenti-PGK-UBE2G1-CR-IRES-Pur, and plenti-PGK-UBE2G1-C90S-CR-IRES-Pur. Gateway donor vectors pDONR223-FLAG-UBE2G1-CR, pDONR223-FLAG-UBE2G1-C90S-CR, pDONR223-FLAG-UBE2D3, and pDONR223-FLAG-UBE2D3-C85S were shuttled into plenti-EF1α-gateway-IRES-Pur via LR recombination to generate plenti-EF1α-FLAG-UBE2G1-CR-IRES-Pur, plenti-EF1α-FLAG-UBE2G1-C90S-CR-IRES-Pur, plenti-EF1α-FLAG-UBE2D3-IRES-Pur, and plenti-EF1α-FLAG-UBE2D3-C85S-IRES-Pur. Constructs pDONR221-U6-sgRNA-EF1a-Cas9-P2A-GFP, plenti-EF1α-Cas9-IRES-Bla, pDONR223-IKZF1, pcDNA3-IKZF1-V5, pcDNA3-CRBN, and pcDNA3-8 x His-Ub were described previously. The Cas9-P2A-GFP coding region of pDONR221-U6-sgRNA-EF1a-Cas9-P2A-GFP was subcloned into pcDNA3.1 (Invitrogen) to generate pcDNA3.1-Cas9-P2A-GFP. The IKZF1 coding region of pDONR223-IKZF1 was shuttle into plenti-EF1α -ePL-gateway-IRES-Bla via LR recombination to generate plenti-EF1α -ePL-IKZF1-IRES-Bla.

Complementary oligonucleotides containing three non-targeting sgRNAs or three gene-specific sgRNAs targeting CRBN or each of the 41 annotated E2 enzymes were annealed and cloned into pRSG16-U6-sgEV-UbiC-TagRFP-2A-Puro or pRSG16-U6-sgEV-UbiC-Hyg, both of which were modified from pRSG16-U6-sg-HTS6C-UbiC-TagRFP-2A-Puro (Cellecta). All sgRNA sequences used in this report (see supplemental table 1) were selected from the human genome-wide CRISPR sgRNA library (Cellecta).

### Lentiviral Production and Transduction

Lentiviral plasmid was cotransfected with the 2nd Generation packaging system (ABM) into 293T cells (Clontech) using Lipofectamine^®^ 2000. After 16 hours of incubation, media was changed to fresh DMEM media supplemented with 20% FBS. At 48 hours post transfection, viral supernatant was collected and cleared via centrifugation at 2000 rpm for 5 minutes, and then filtered through a 0.45 micron cellulose acetate or nylon filter unit. Acute myeloid leukemia and myeloma cell lines were spin-inoculated with lentivirus at 2500 rpm for 120 minutes. After twelve hours, viral supernatant was removed and complete culture media was added to the cells. Forty-eight hours later, cells were incubated with 1~2 μg/mL puromycin (Thermofisher), 10~20 μg/mL blasticidin (Thermofisher), or 250~500 μg/mL hygromycin B (Thermofisher) for an additional 2~7 days to select cells stably integrated with lentiviral vectors.

### CRISPR Gene Editing

AML and MM cell lines were transduced with plenti-EF1a-Cas9-IRES-Bla, followed by limiting dilution and blasticidin selection in 96-well plates (Corining) to generate single clones stably expressing Cas9. The expression of Cas9 in stable clones were validated by immunoblot analysis. Next, Cas9-expressing cells were transduced with pRSG16-U6-sgNT-1-UbiC-TagRFP-2A-Puro, pRSG16-U6-sgNT-3-UbiC-TagRFP-2A-Puro, pRSG16-U6-sgUBE2G1-1-UbiC-TagRFP-2A-Puro, pRSG16-U6-sgUBE2G1-5-UbiC-TagRFP-2A-Puro, pRSG16-U6-sgUBE2D3-4-UbiC-TagRFP-2A-Puro, or pRSG16-U6-sgCRBN-8-UbiC-TagRFP-2A-Puro. One days after transduction, cells were selection with puromycin for 2 days. Then, gene editing efficiency was verified by immunoblot analysis with antibodies recognizing the targeted proteins. For gene editing of both UBE2G1 and UBE2D3, U937-Cas9 cells were first transduced with pRSG16-U6-sgUBE2G1-5-UbiC-Hyg. After 24 hours, cells were selected with hygromycin B selection for additional 5 days, and then transduced with pRSG16-U6-sgUBE2D3-4-UbiC-TagRFP-2A-Puro, followed by puromycin selection for 2 days and immunoblot analysis. 293T cells were transiently transfected with pcDNA3.1-Cas9-P2A-GFP, pRSG16-U6-sgUBE2G1-5-UbiC-TagRFP-2A-Puro with or without pRSG16-U6-sgUBE2D3-4-UbiC-TagRFP-2A-Puro. Three days after transfection, cells were subjected to limiting dilution into 96-well plates. After two weeks, stable clones were cherry-picked, expanded and subjected to immunoblot analysis. 293T *UBE2G1-/-* clone 13 was validated to be UBE2G1 deficient, and *UBE2G1-/-;UBE2D3-/-* clone 4 was proven deficient for both UBE2G1 and UBE2D3.

For the single-guide RNA directed E2 CRISPR screen, U937 Cas9 cells were first transduced with plenti-EF1a-ePL-IKZF1-IRES-Bla to generate U937_Cas9_ePL-IKZF1 cells, which were then transduced with the focused lentiviral CRISPR library containing 3 non-targeting control sgRNAs and 3 gene-specific sgRNAs targeting each of the 41 E2 enzymes.

For the dual-guide RNA directed E2 CRISPR screen, U937_Cas9_ePL-IKZF1 were transduced with pRSG16-U6-sgUBE2G1-5-UbiC-Hyg, followed by hygromycin B selection for additional 5 days. Then, cells were transduced with the focused lentiviral CRISPR library targeting all annotated E2s as described above.

### IKZF1-ePL Degradation Assay

Four days after transduction with lentiviral vectors expressing non-targeting, *CRBN*-specific or *UBE2*-specific sgRNA, U937_Cas9_ePL-IKZF1 cells were dispensed into a 384-well plate (Corning) pre-spotted with pomalidomide at varying concentrations. Twenty-five microliters of RPMI-1640 growth media containing 5000 cells was dispensed into each well. After incubation at 37°C with 5% CO2 for 16 hrs, 25 μL of the InCELL Hunter™ Detection Reagent Working Solution (DiscoverX) was added to each well and incubated at RT for 30 minutes protected from light. After 30 minutes, luminescence was read on an EnVision Multimode Plate Reader (Perkin Elmer). For each cell line, The IKZF1 degradation induced by pomalidomide at each indicated concentration was normalized with the DMSO control. Then, a four parameter logistic model (sigmoidal dose-response model) was used to plot the IKZF1 destruction curves. All percentage of control IKZF1 destruction curves were processed and graphed using GraphPad Prism,Version 7.

### Cell Proliferation Assay

AML cell lines (5000 cells per well), MM cell lines (5000 cells per well) or 293T cells (2000 cells per well) in 50 μL complete culture media were seeded into black 384-well plates containing with DMSO or test compounds. After 3 or 5 days, cell proliferation was assessed using CTG according to manufacturer’s instructions. Relative cell proliferation was normalized against the DMSO control. The growth inhibitory curve of each test compound was processed and graphed using GraphPad Prism,Version 7.

### Immunoblot Analysis

Following treatment with test compounds at 37°C for the indicated time, cells were washed in ice-cold 1X PBS twice before harvest in Buffer A [50 mM Tris.Cl (pH 7.6), 150 mM NaCl, 1% Triton X-100, 1 mM EDTA, 1 mM EGTA, 1 mM β-glycerophosphate, 2.5 mM sodium pyrophosphate, 1 mM Na3VO4, 1 μg/mL leupeptin, one tablet of Complete ULTRA protease inhibitor cocktail (Roche), and one tablet of PhosSTOP phosphatase inhibitor cocktail (Roche)]. Whole cell extracts were collected after centrifugation at top speed for 10 minutes, resolved by SDS-PAGE gel electrophoresis, transferred onto a nitrocellulose membrane using the Turboblot system (Bio-Rad), and probed with the indicated primary antibodies. Bound antibodies were detected with IRDye-680 or −800 conjugated secondary antibodies using a LI-COR scanner.

### Protein Half Life Analysis

293T parental and *UBE2G1-/-* cells were pretreated with DMSO or pomalidomide (1 μM) for 30 minutes, followed by the addition of 100 μg/ml cycloheximide (EMD) into the culture medium. At various time points as indicated in Fig. 3D, cells were collected and subjected to immunoblot analysis.

### *In Vivo* Ubiquitination Assay

The ubiquitination assays were carried out as described previously (Lu et al., 2014b). In brief, 293T parental, *UBE2G1-/-*, and *UBE2G1-/-;UBE2D3-/-* cells seeded in 6 well plates were transiently transfected with pcDNA3-IKZF1-V5, pcDNA3-CRBN, pcDNA3-8 x His-Ub, or pcDNA3 empty vector. In Figures 5B and S6B-D, 293T *UBE2G1-/-* and *UBE2G1-/-;UBE2D3-/-* cells were also transfected with plenti-EF1a-UBE2G1-IRES-Pur, plenti-EF1a-UBE2D3-IRES-Pur or both. Forty-eight hours post transfection, cells were treated with 10 μM MG132 with or without an increased concentrations of pomalidomide as indicated. Eight hours later, cells were washed twice with ice cold PBS and resuspended in 1 mL PBS. Twenty uL of the cell suspension was boiled in LDS loading buffer, and the remaining cells were collected via centrifugation and lysed in Buffer C (6M guanidine-HCL, 0.1M Na2HPO4/NaH2PO4, 20 mM imidazole, pH 8.0). Next, whole cell extracts were sonicated for 12 pulses, and mixed with 20 μL of HisPur™ Ni-NTA Magnetic Beads (Thermofisher) at 37°C for 4 hours. Ni-NTA beads were then washed three times with Buffer C, three times with Buffer D (1 Volume of Buffer C: 3 volumes of Buffer E), and three times with Buffer E (25 mM Tris.CL, 20 mM imidazole, pH 6.8). Bound proteins were eluted by boiling in 2x LDS loading buffer and subjected to immunoblot analysis.

### Antibodies

Rabbit anti-human CRBN65 monoclonal antibody (mAb) (Celgene, San Diego, CA); rabbit anti-human GSPT1 polyclonal antibody (pAb; Abcam), rabbit anti-human IKZF1 mAb (Cell Signaling), rabbit anti-human IKZF3 mAb (Cell Signaling), rabbit anti-human CK1α pAb (Abcam), rabbit anti-human ZFP91 pAb (LifeSpan Biosciences), mouse anti-human UBE2G1 mAb (Santa Cruz), rabbit anti-human UBE2D3 pAb (Sigma), rabbit anti-human Cul4A pAb (Cell Signaling), rabbit anti-human DDB1 pAb (Cell Signaling), rabbit anti-human Rbx1 pAb (Cell Signaling), rabbit anti-human Cdt1 pAb (Cell Signaling), rabbit anti-human Cdt2 pAb (Cell Signaling), rabbit anti-human Set8 pAb (Cell Signaling), rabbit anti-human RBM39 pAb (Sigma), rabbit anti-human p21 pAb (Cell Signaling), rabbit anti-human p27 pAb (Cell Signaling), rabbit anti-human c-Myc pAb (Cell Signaling), mouse anti-penta-HIS mAb (Qiagen), and mouse anti-human Actin and Tubulin antibodies (Sigma) were used as primary antibodies. Goat anti-mouse 800 antibody (LI-COR Biosciences), goat anti-rabbit 680 antibody (LI-COR Biosciences), goat anti-mouse 800 antibody (LI-COR Biosciences) and goat anti-rabbit 680 antibody (LI-COR Biosciences) were used as secondary antibodies.

---

**Figure S1.**
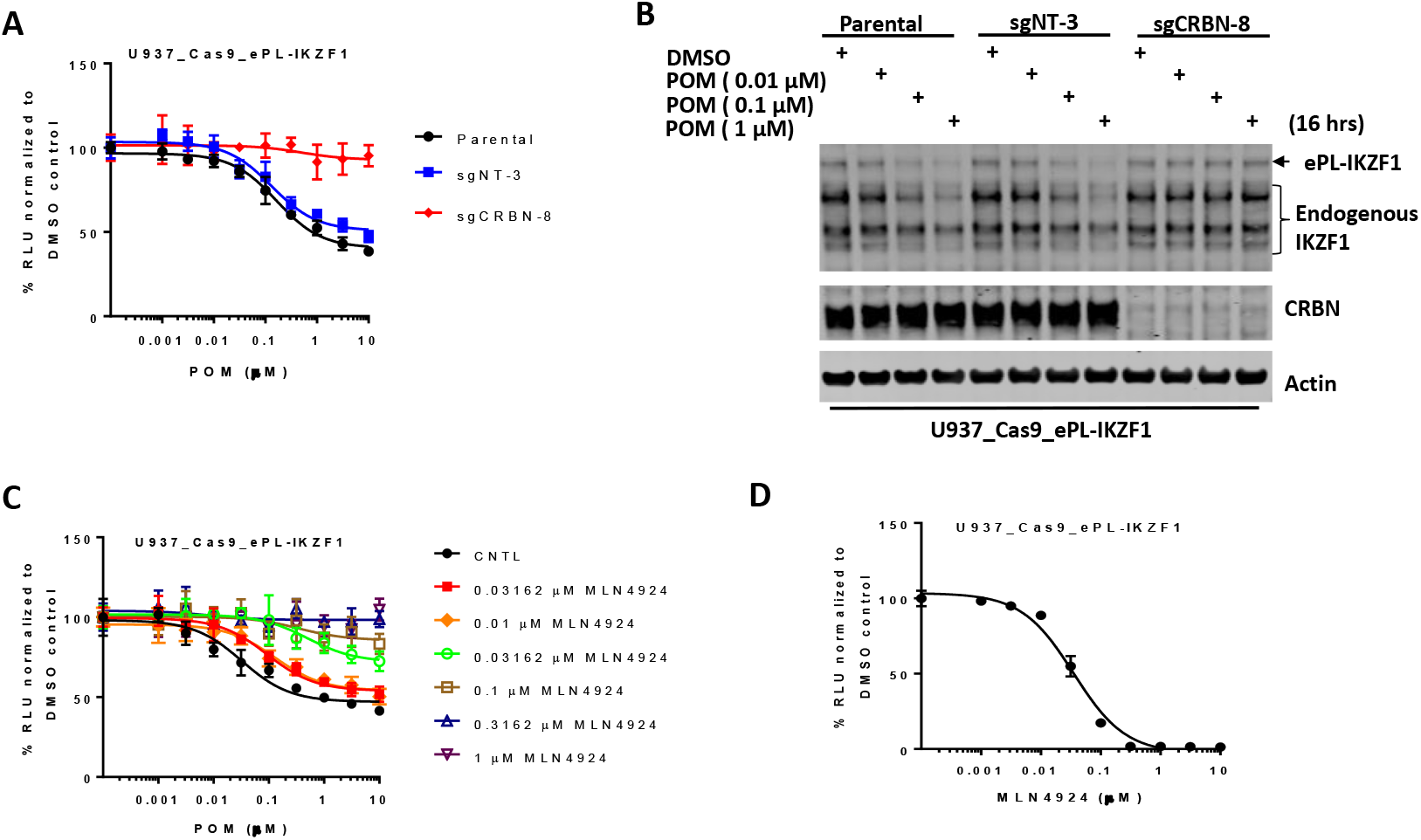
Pomalidomide-induced destruction of IKZF1 requires CRL4^CRBN^ (A and B) Chemiluminescent measurement (A) or immunoblot analysis (B) of ePL-IKZF1 protein expression level in U937_Cas9_ePL-IKZF1 parental cells or cells expressing non-targeting or *CRBN*-specific sgRNA. Cells were treated with DMSO or an increasing concentrations of POM for 16 hours. In (A), data are presented as mean ± SD (n = 4). In (B), note that the degradation efficiency of ePL-tagged and endogenous IKZF1 is comparable. (C) Chemiluminescent measurement of ePL-IKZF1 protein expression level in U937_Cas9_ePL-IKZF1 cells treated with DMSO or an increasing concentrations of POM in the presence or absence or MLN4924 at the indicated concentrations for 16 hours. Data are presented as mean ± SD (n=4). (D) Cell proliferation of U937_Cas9_ePL-IKZF1 cells treated with DMSO or MLN4924 at the indicated concentrations for 48 hours. Cell proliferation was determined by CTG. Data are presented as mean ± SD (n=3).

**Figure S2.**
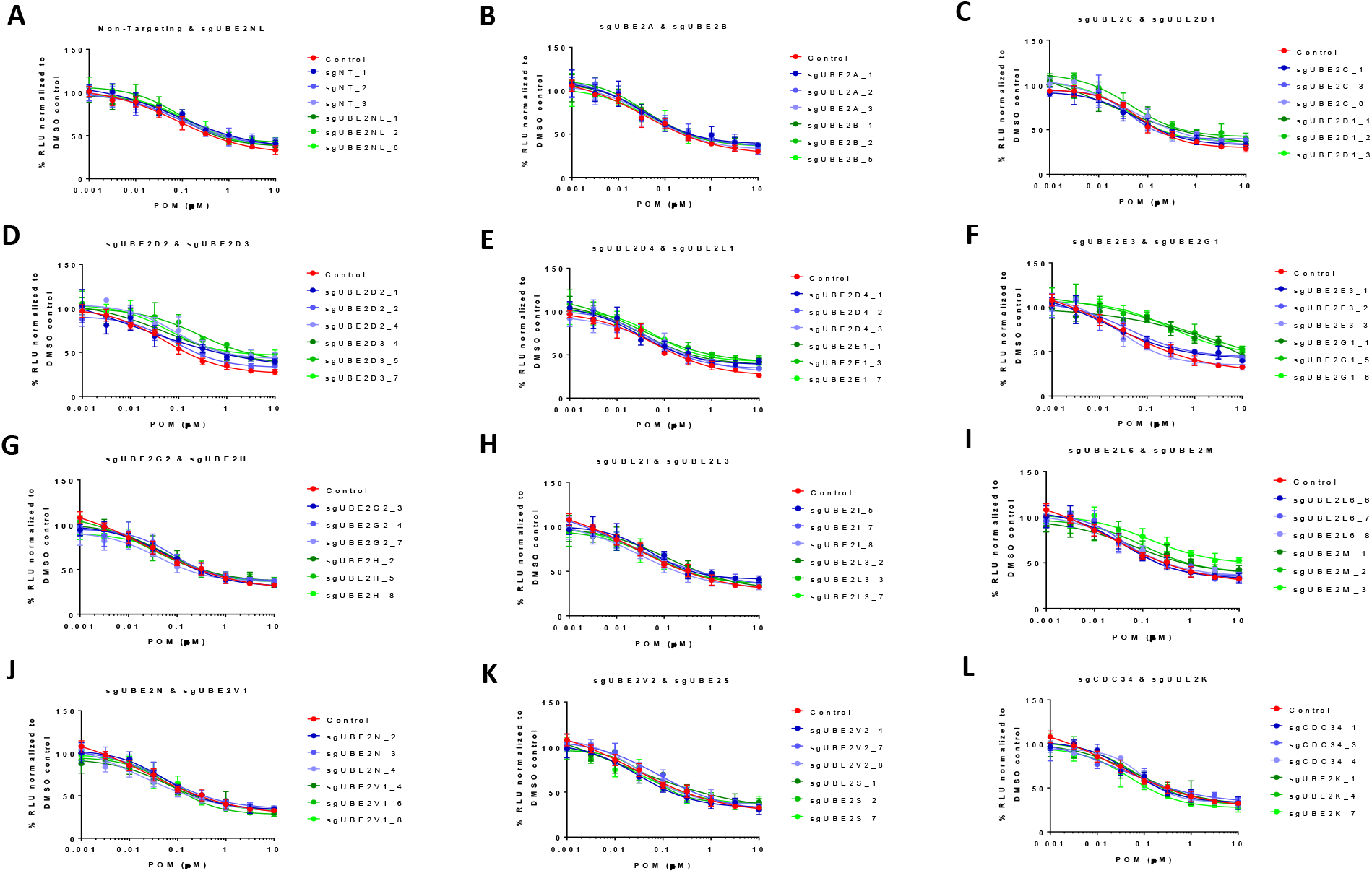
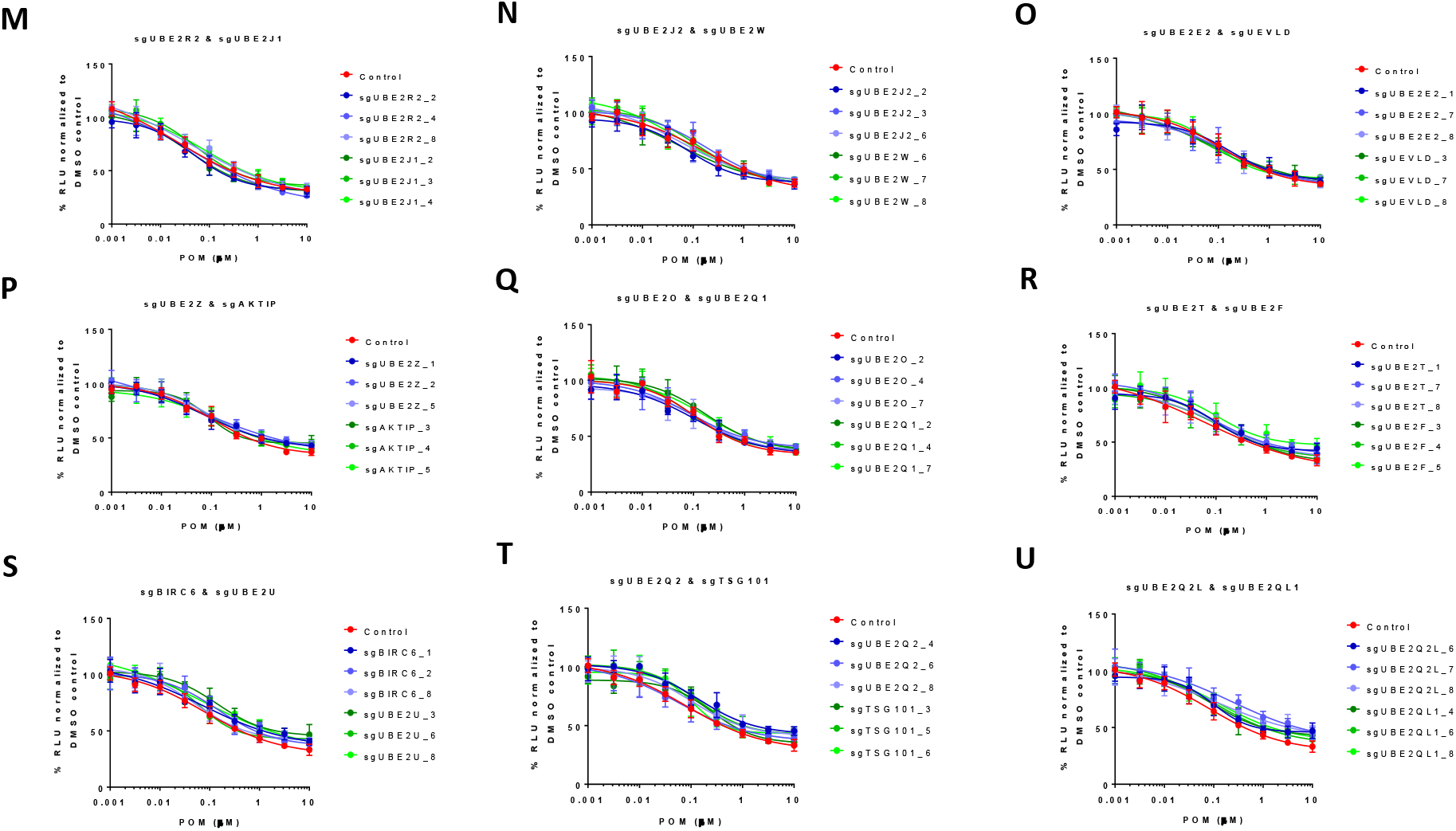
The effect of individual knockout of each of the 41 annotated E2 enzymes on pomalidomide-induced destruction of IKZF1 (A-U) Chemiluminescent measurement of ePL-IKZF1 protein expression level in U937_Cas9_ePL-IKZF1 parental cells or cells expressing non-targeting or E2-specific sgRNA. Cells were treated with DMSO or an increasing concentrations of POM for 16 hours. Data are presented as mean ± SD (n = 4).

**Figure S3.**
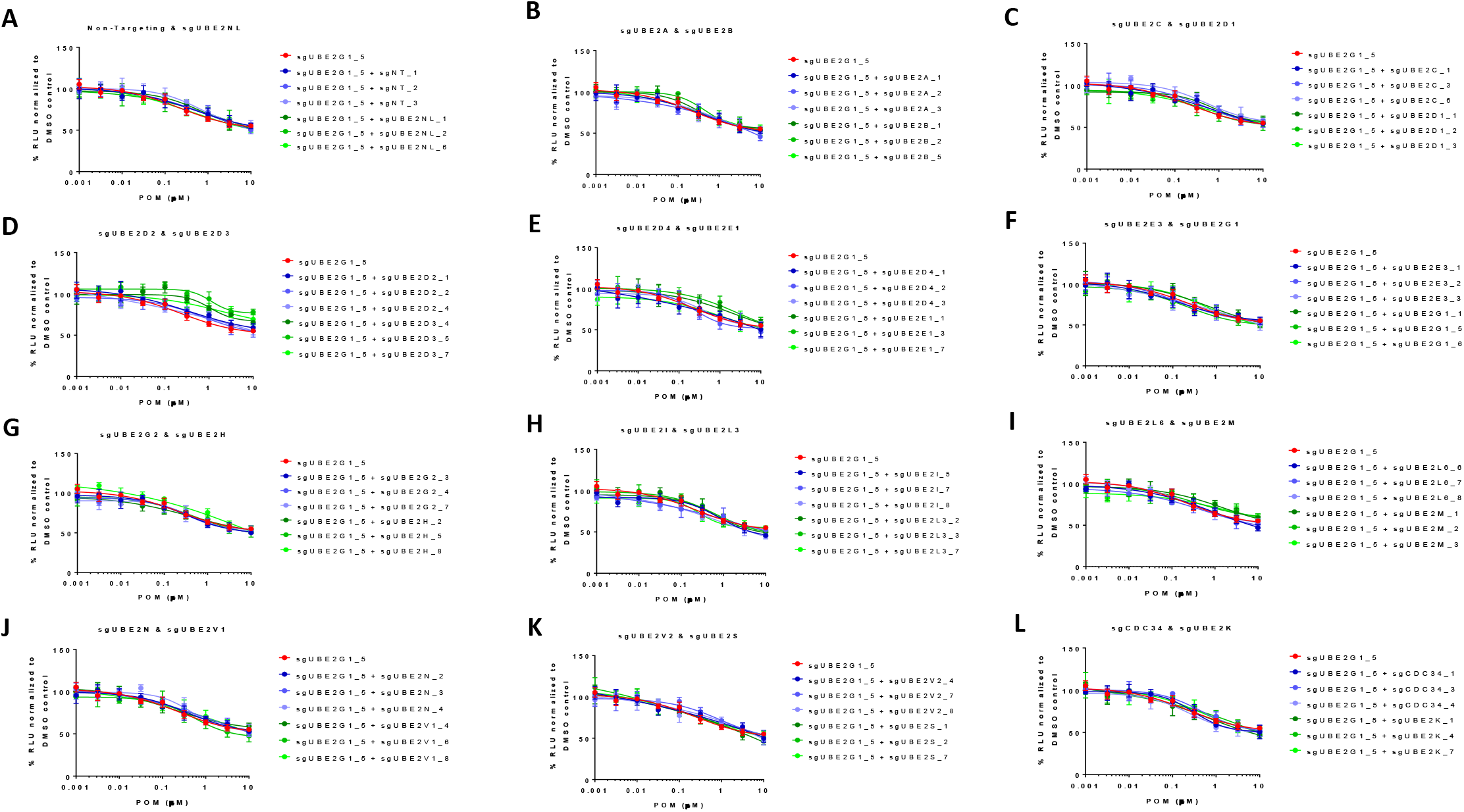
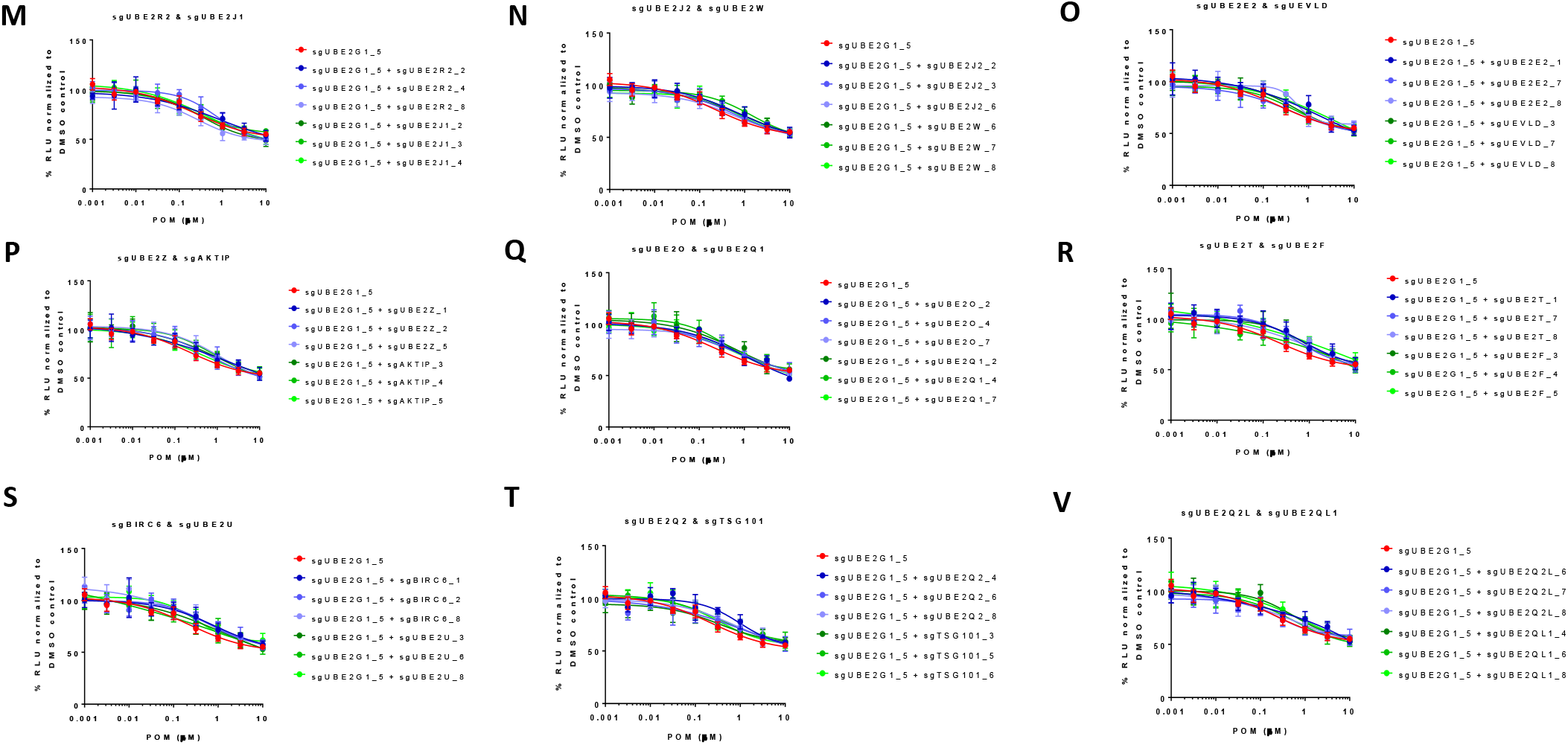
The effect of double knockout of UBE2G1 and one of the 41 E2 enzymes on pomalidomide-induced destruction of IKZF1 (A-U) Chemiluminescent measurement of ePL-IKZF1 protein expression level in U937_Cas9_ePL-IKZF1 parental cells or cells expressing both UBE2G1-specific and non-targeting or E2-specific sgRNA. Cells were treated with DMSO or an increasing concentrations of POM for 16 hours. Data are presented as mean ± SD (n = 4).

**Figure S4.**
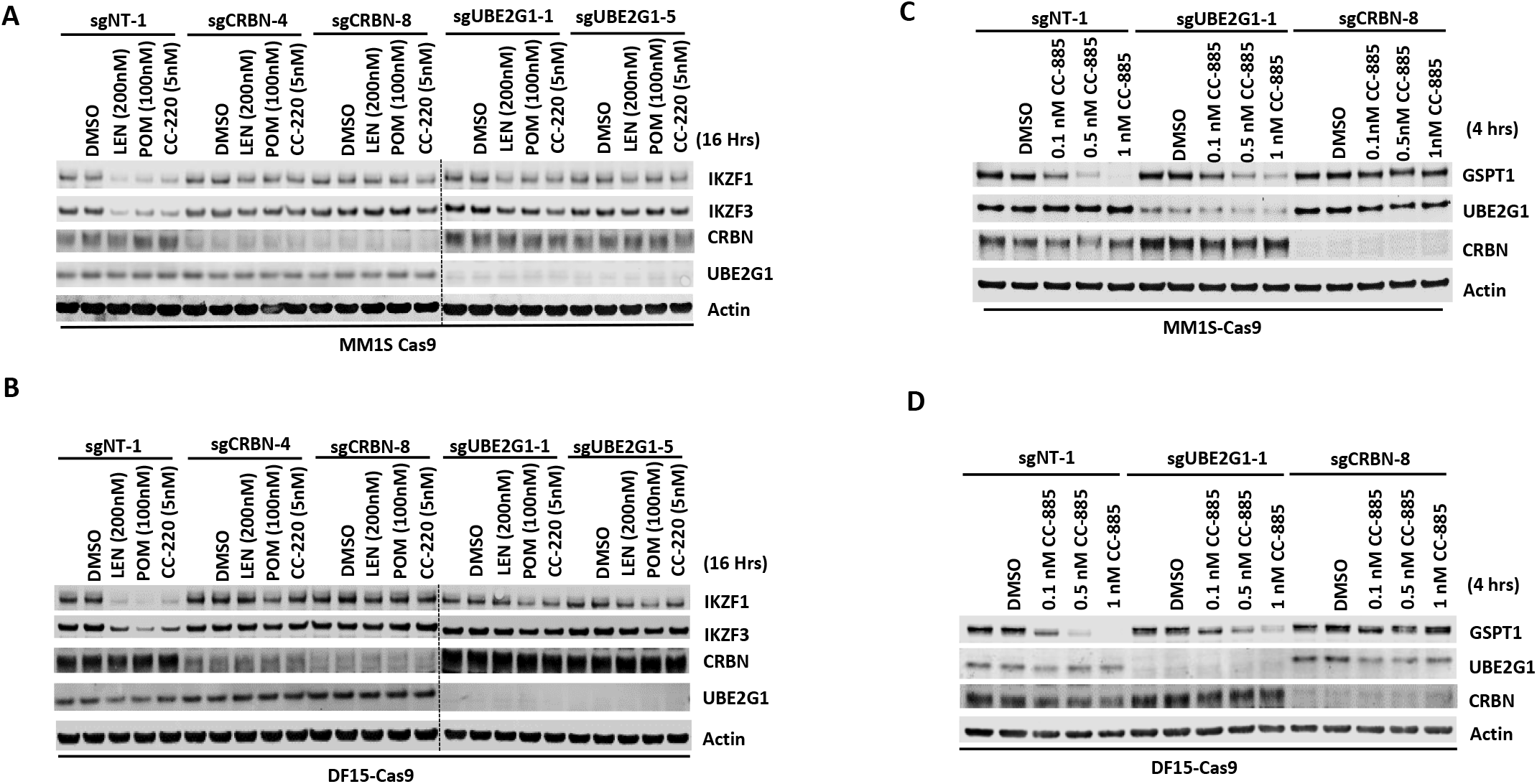
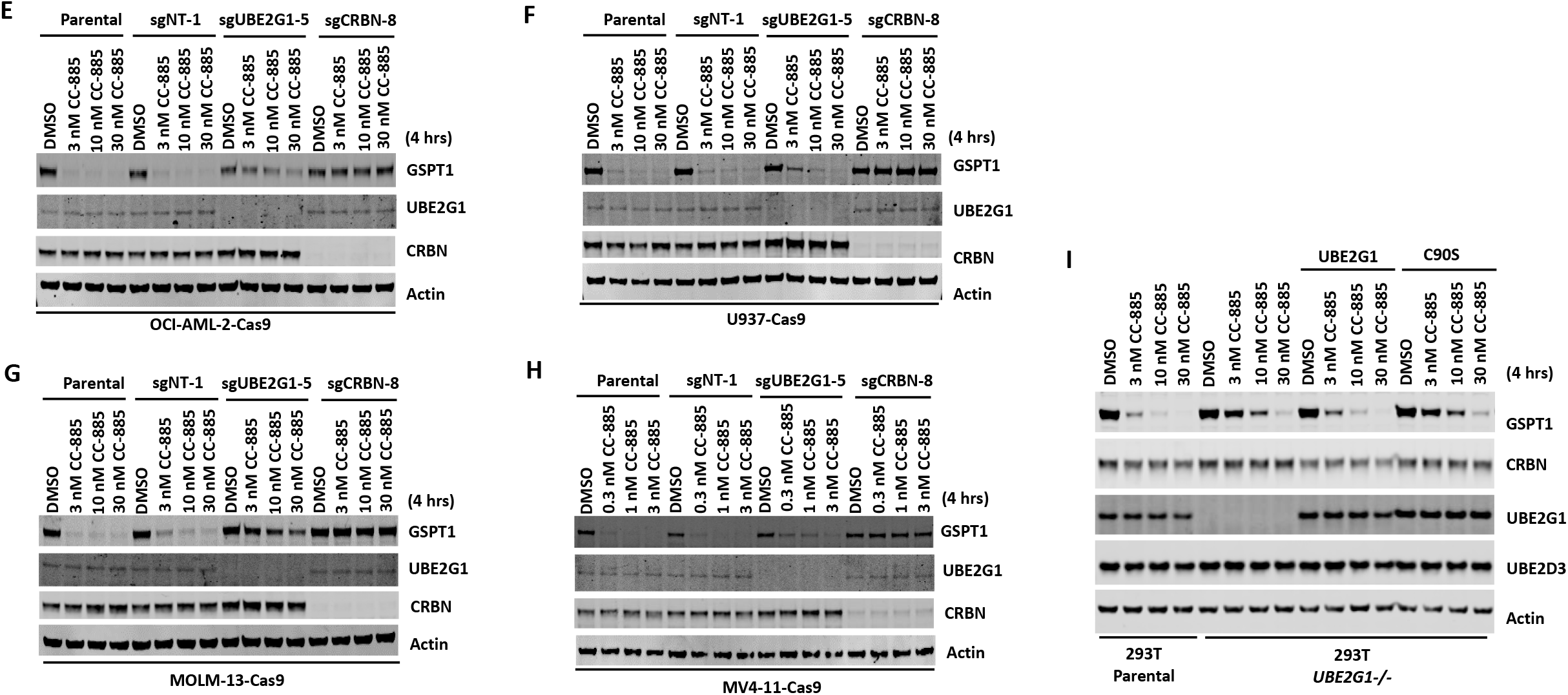
Elimination of UBE2G1 blocks the degradation of cereblon neomorphic substrates recruited by lenalidomide and CC-885 (A-F) Immunoblot analysis of DF15-Cas9 cells (A and C), MM1S-Cas9 cells (B and D), OCI-AML2-Cas9 cells (E), U937-Cas9 cells (F), MOLM-13-Cas9 cells (G) and MV4-11-Cas9 cells (H) transduced with lentiviral vectors expressing non-targeting, UBE2G1-specific or *CRBN*-specific sgRNAs. Cells were treated with lenalidomide for 16 hours (A and B), or CC-885 for 4 hours (C-H) at the indicated concentrations. (I) Immunoblot analysis of 293T parental or *UBE2G1-/-* cells stably expressing UBE2G1 wild-type or C90S mutant. Cells were treated with CC-885 at the indicated concentrations for 4 hours.

**Figure S5.**
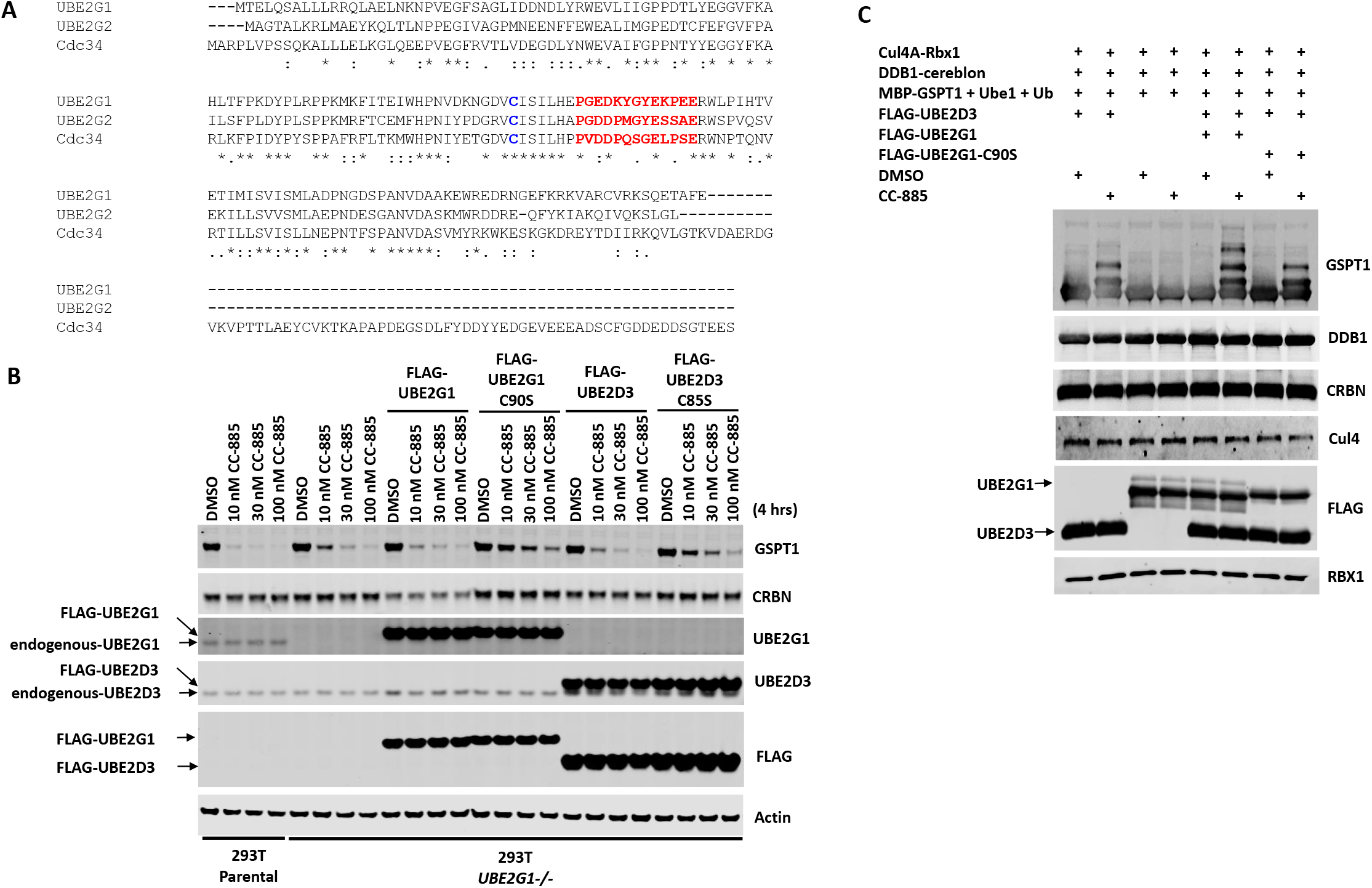
UBE2G1 catalyzes the ubiquitin chain assembly on GSPT1 pre-conjugated with ubiquitin (A) Sequence alignment of human UBE2G1, human UBE2G2 and human CDC34 using Clustal W 2.1. The acidic loops indispensable for the assembly of K48-linked ubiquitin chains are highlighted with red, and the catalytic cysteines are highlighted with blue. (B) Immunoblot analysis of 293T parental or *UBE2G1-/-* cells transduced with lentiviral vectors expressing FLAG-tagged UBE2G1 wild-type or C90S mutant, or FLAG-tagged UBE2D3 wild-type or C85S mutant. Cells were treated with CC-885 at the indicated concentrations for 4 hours. Note that overexpression of wild-type FLAG-UBE2G1 or FLAG-UBE2D3 partially rescued the GSPT1 degradation defect caused by UBE2G1 deficiency, while overexpression of catalytically-dead mutant FLAG-UBE2G1-C90S or FLAG-UBE2D3-C85S further blocked the degradation of GSPT1. (C) *In vitro* ubiquitination of GSPT1 by CRL4^CRBN^ with or without CC-885 and indicated E2 variants. Consistent with results observed with bacterial recombinant UBE2G1 and UBE2D3 proteins, FLAG-UBE2G1 and FLAG-UBE2D3 proteins purified from human cells acted in concert to promote the ubiquitination of GSPT1.

**Figure S6.**
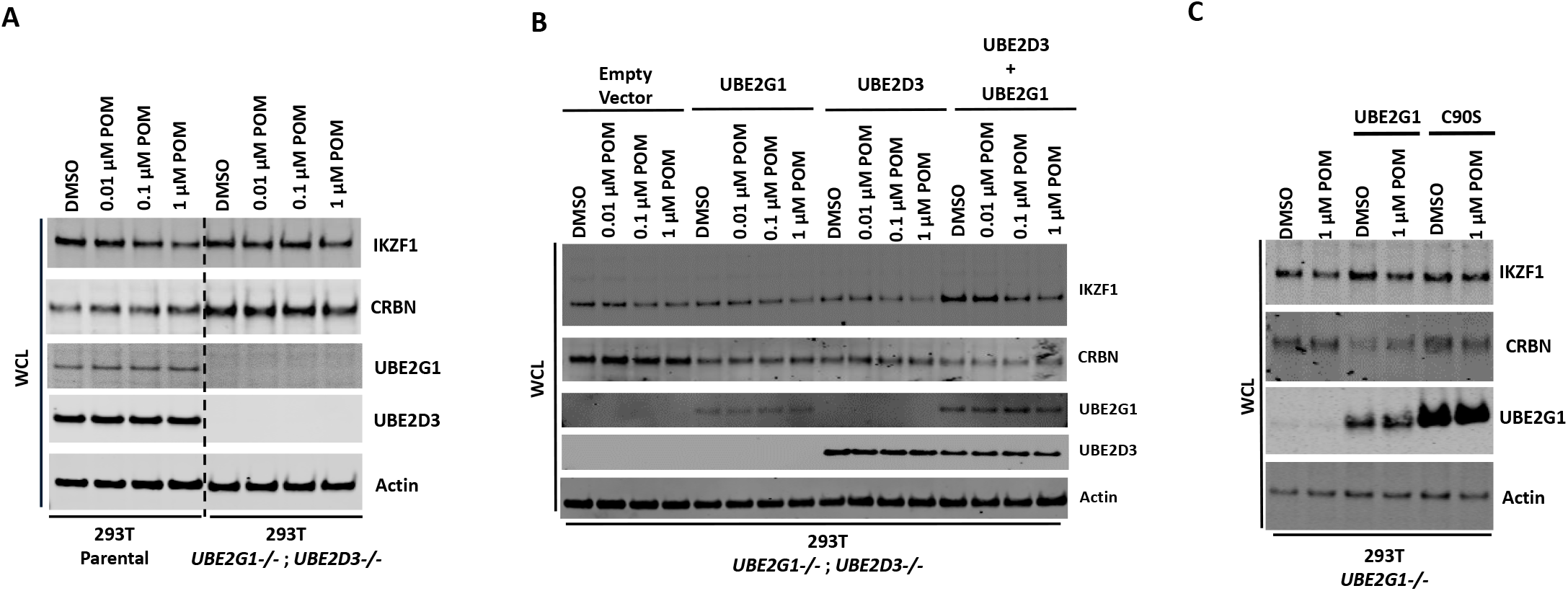
Input protein levels for the *in vivo* ubiquitinaiton studies corresponding to Figure 5 (A) Total input for Figures 5A. Immunoblot analysis of 293T parental and *UBE2G1-/- ; UBE2D3-/-* (Clone 4) cells transfected to produce 8xHis-Ubiquitin, cereblon and IKZF1-V5. (B) Total input for Figure 5B. Immunoblot analysis of 293T parental and *UBE2G1-/-;UBE2D3-/-* (Clone 4) cells transfected to produce 8xHis-Ubiquitin, CRBN, IKZF1-V5 with or without UBE2G1 and/or UBE2D3. (C) Total input for Figure 5C. Immunoblot analysis of 293T parental and *UBE2G1-/-* (Clone 13) cells transfected to produce 8xHis-Ubiquitin, CRBN, IKZF1-V5 with or without UBE2G1 wildtype or C90S mutant. In (A), (B) or (C), 48 hours after transfection, cells were treated with 10 μM MG132 and POM at the indicated concentrations for additional 8 hours.

**Figure S7.**
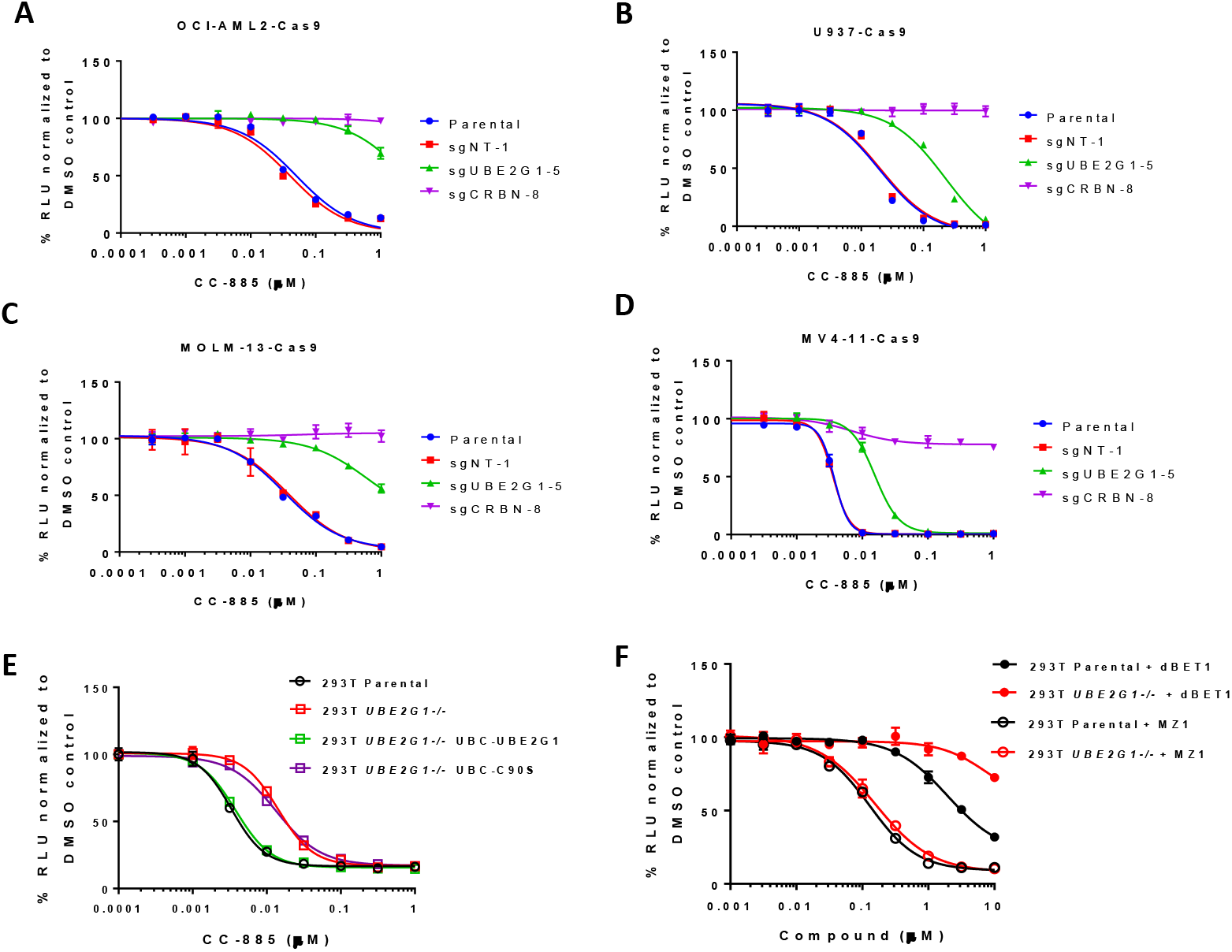
The growth-inhibitory effect of CC-885 and Brd4 PROTACs in AML cell lines and 293T cells (A-D) Cell proliferation of AML cell lines OCI-AML2 (A), U937 (B), MOLM-13 (C) and MV4-11 (D) treated with DMSO or CC-885 at the indicated concentrations for 72 hours. Cell proliferation was determined by CTG. Data are presented as mean ± SD (n=4). (E and F) Cell proliferation of 293T parental and *UBE2G1-/-* (clone 13) cells with or without ectopic overexpression of UBE2G1 wild-type or C90S mutant. Cells were treated with CC-885 (E), dBET1 (F) or MZ-1 (F). Cell proliferation was determined by CTG. Data are presented as mean ± SD (n=4).

**Figure S8.**
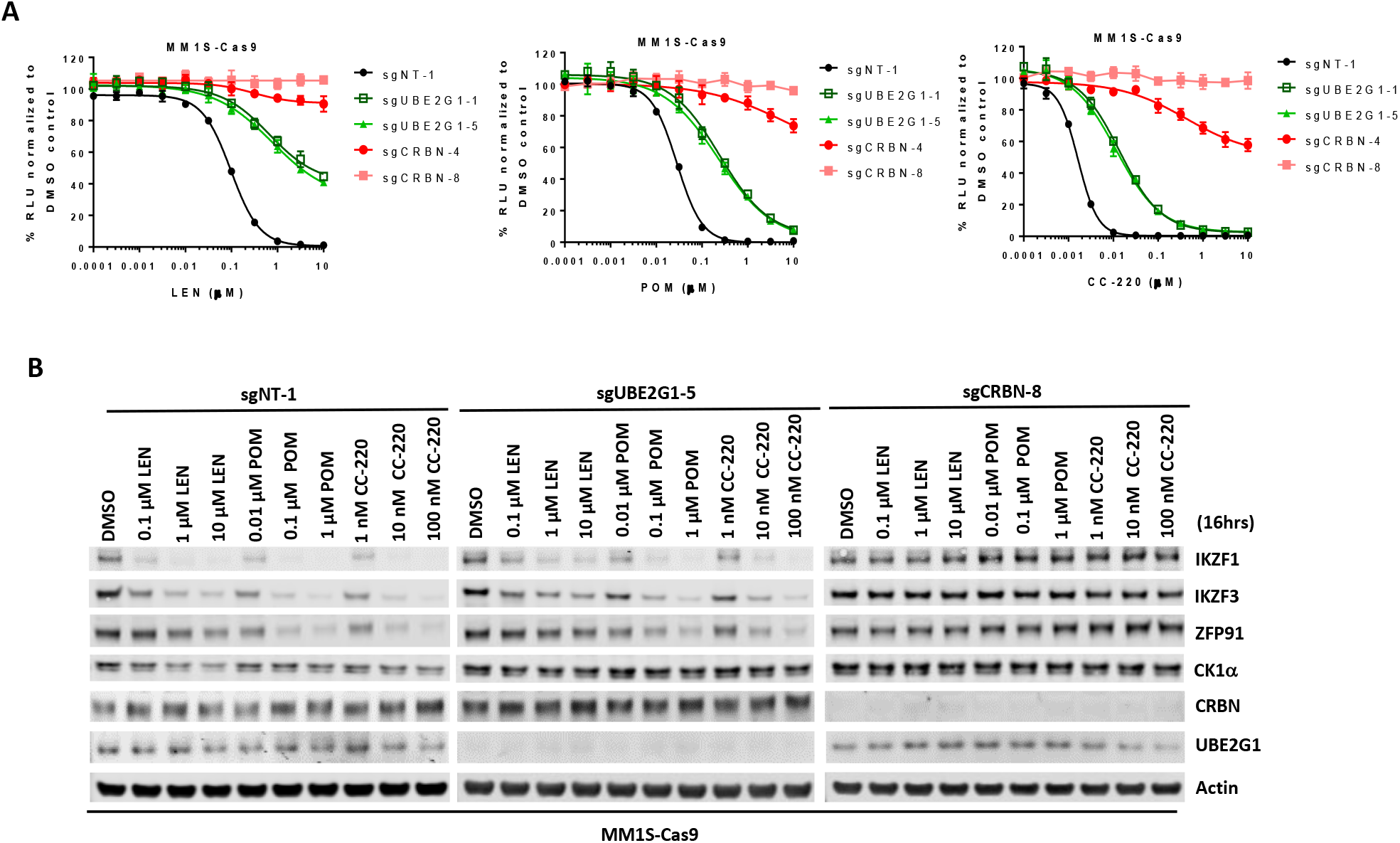
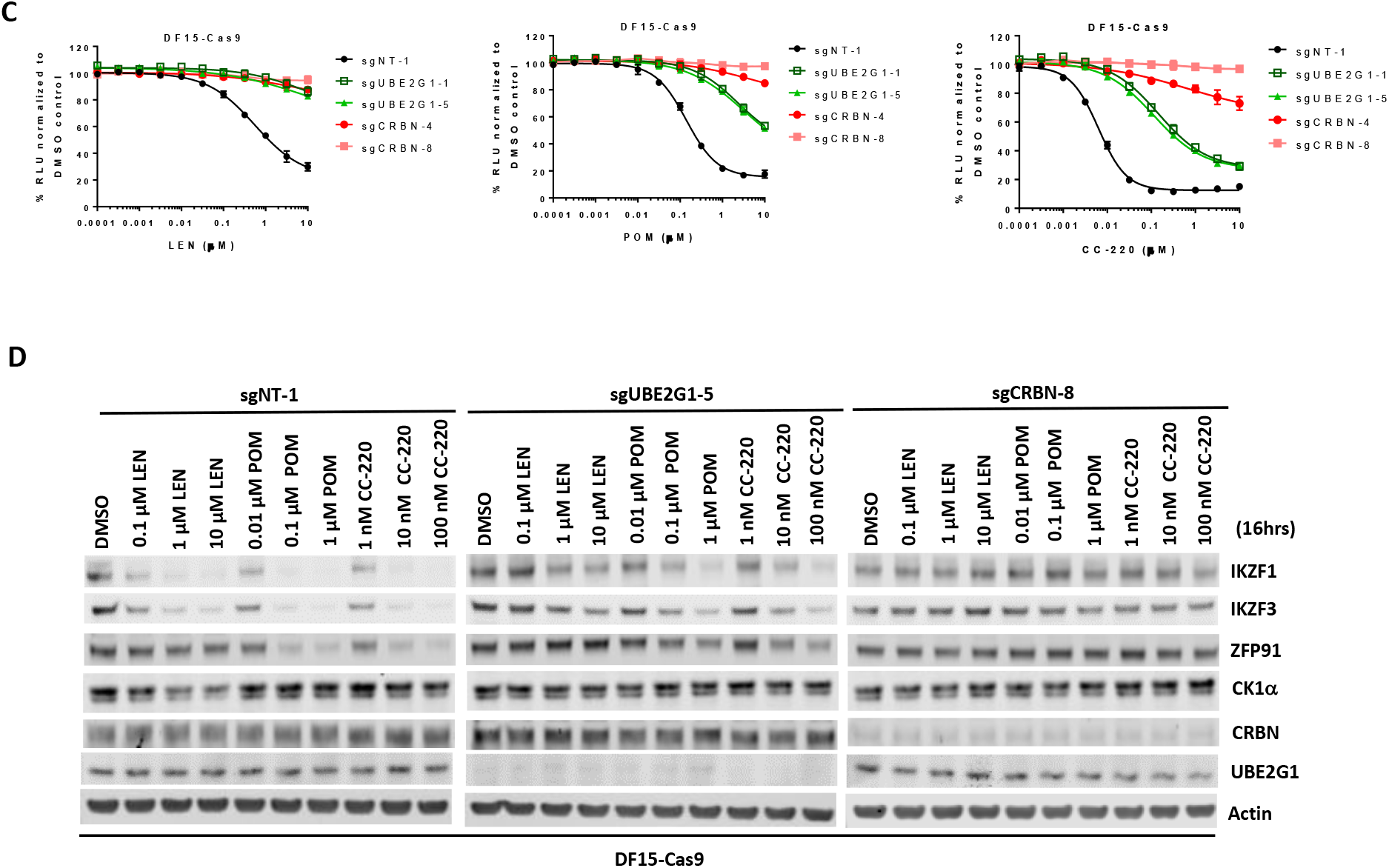
UBE2G1 knockout diminished the responses to lenalidomide, pomalidomide and CC-220 in myeloma cell lines DF15 and MM1S (A-D) Cell proliferation (A and C) and immunoblot analysis (B and D) of MM1S-Cas9 (A and B) and DF15-Cas9 (C and D) cells infected with lentiviral vectors expressing non-targeting, UBE2G1-specific or CRBN-specific sgRNAs. Cells were treated DMSO vehicle control, LEN, POM or CC-220 at the indicated concentrations for 5 days (A and C) or 16 hours (B and D). In (A and C), cell proliferation was determined by CTG, and data are presented as mean ± SD (n=3).

**Figure S9.**
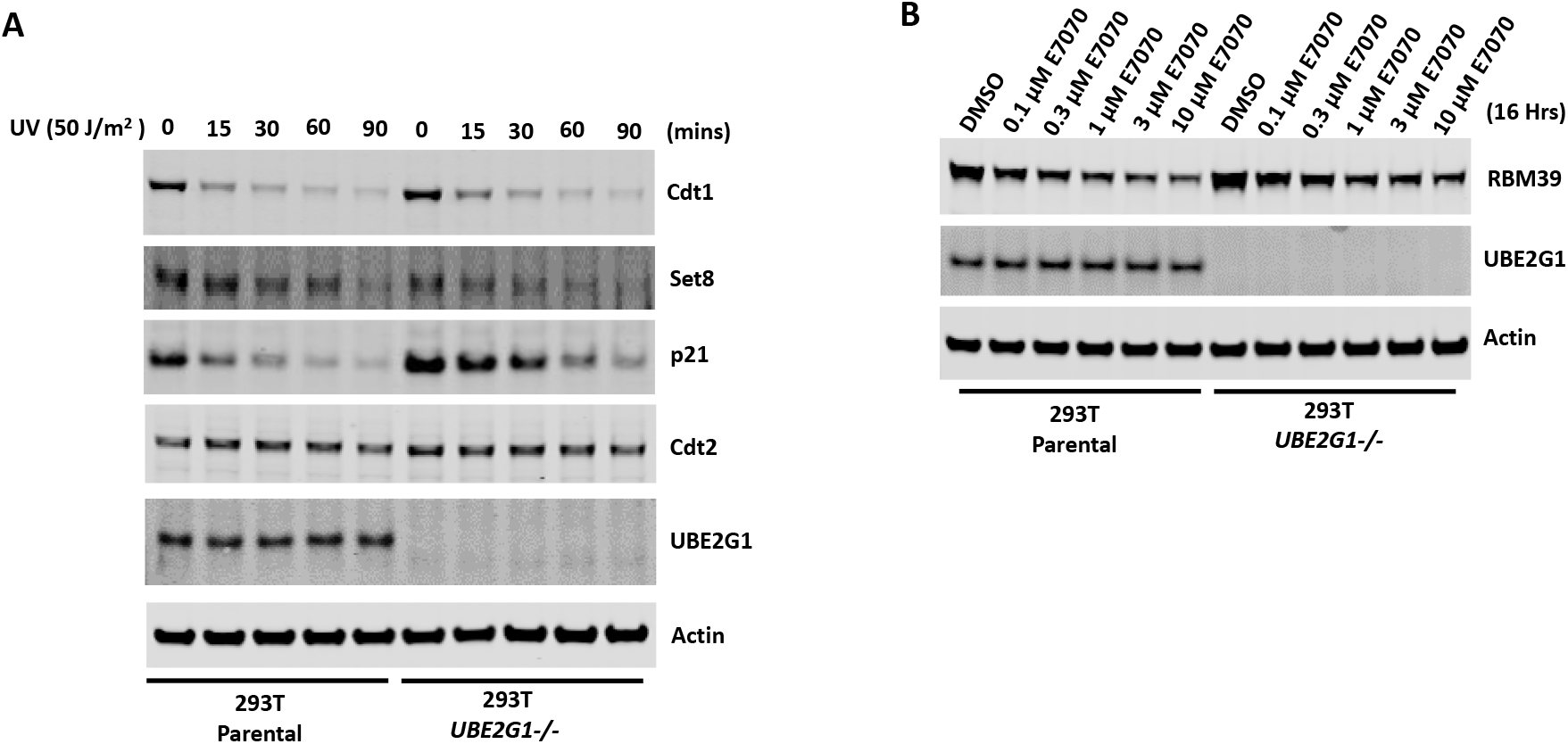
Depletion of UBE2G1 attenuated the degradation of p21 and RMB39 induced by UV irradiation and sulfonamide treatment, respectively. (A and B) Immunoblot analysis of 293T parental and UBE2G1-/- (clone 13) cells treated with UV irradiation (A) or E7070 (B). In (A), cells were UV irradiated at 50 J/m2 using a Stratalinker, and collected at the indicated time points thereafter. In (B), cells were treated with DMSO or an increasing concentration of E7070 for 16 hours.

**Figure S10.**
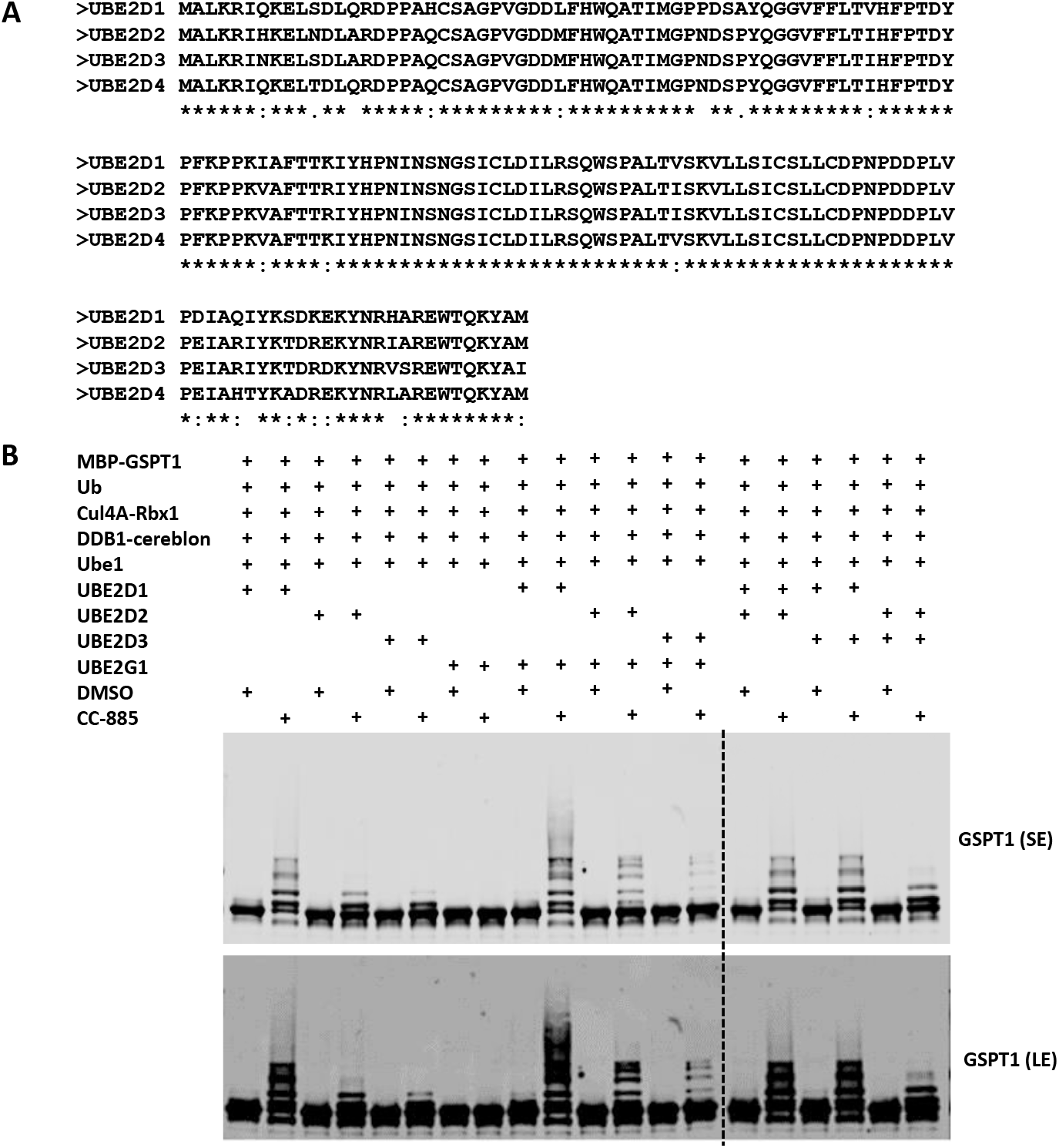
UBE2D family proteins redundantly promote the ubiquitination of GSPT1 (A) Sequence alignment of human UBE2D family proteins using Clustal W 2.1. Note that the amino acid sequence identity among all 4 family proteins is close to 90%. (B) In vitro ubiquitination of GSPT1 MBP fusion protein by recombinant CRL4CRBN complex in the presence of UBE2G1, UBE2D1, UBE2D2, or UBE2D3, alone or in combination. Recombinant protein products as indicated were incubated with or without 80 μM CC-885 in the ubiquitination assay buffer at 30 °C for 2 hours, and then analyzed by immunoblotting. SE, short exposure; LE, long exposure

**Supplemental Table 1.**
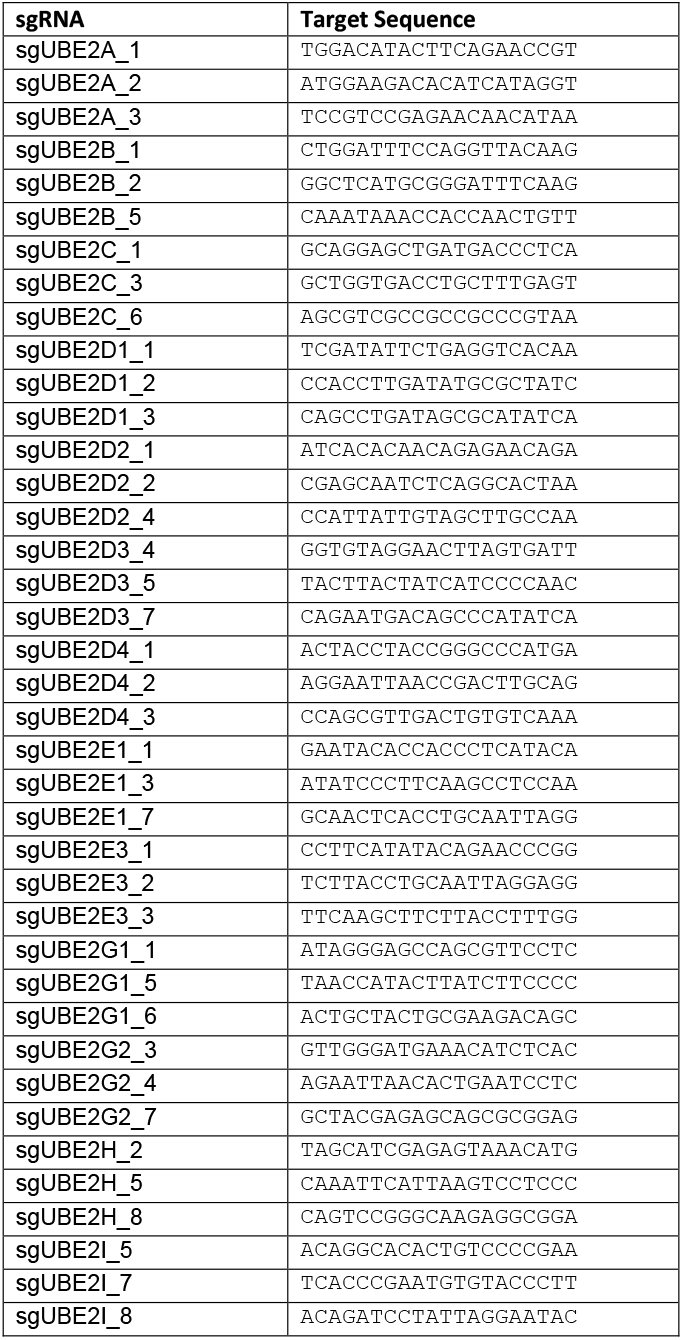
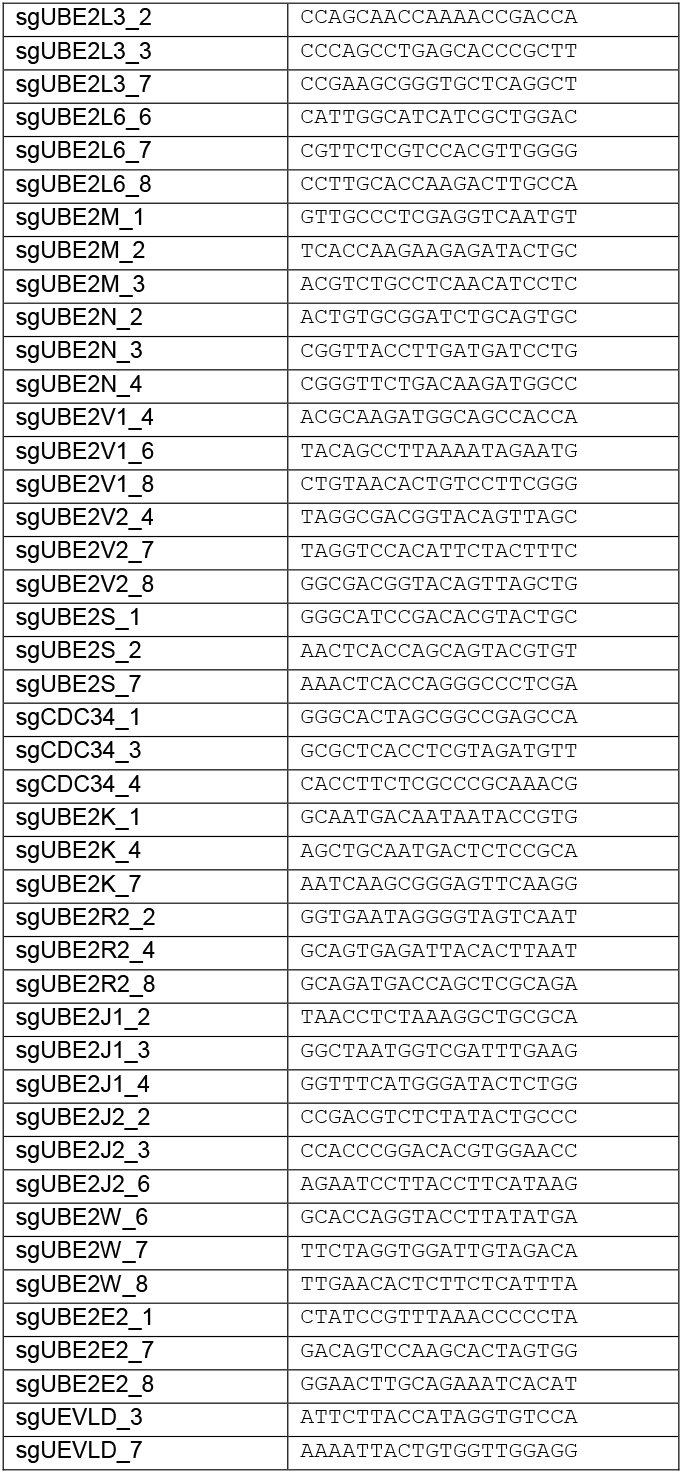
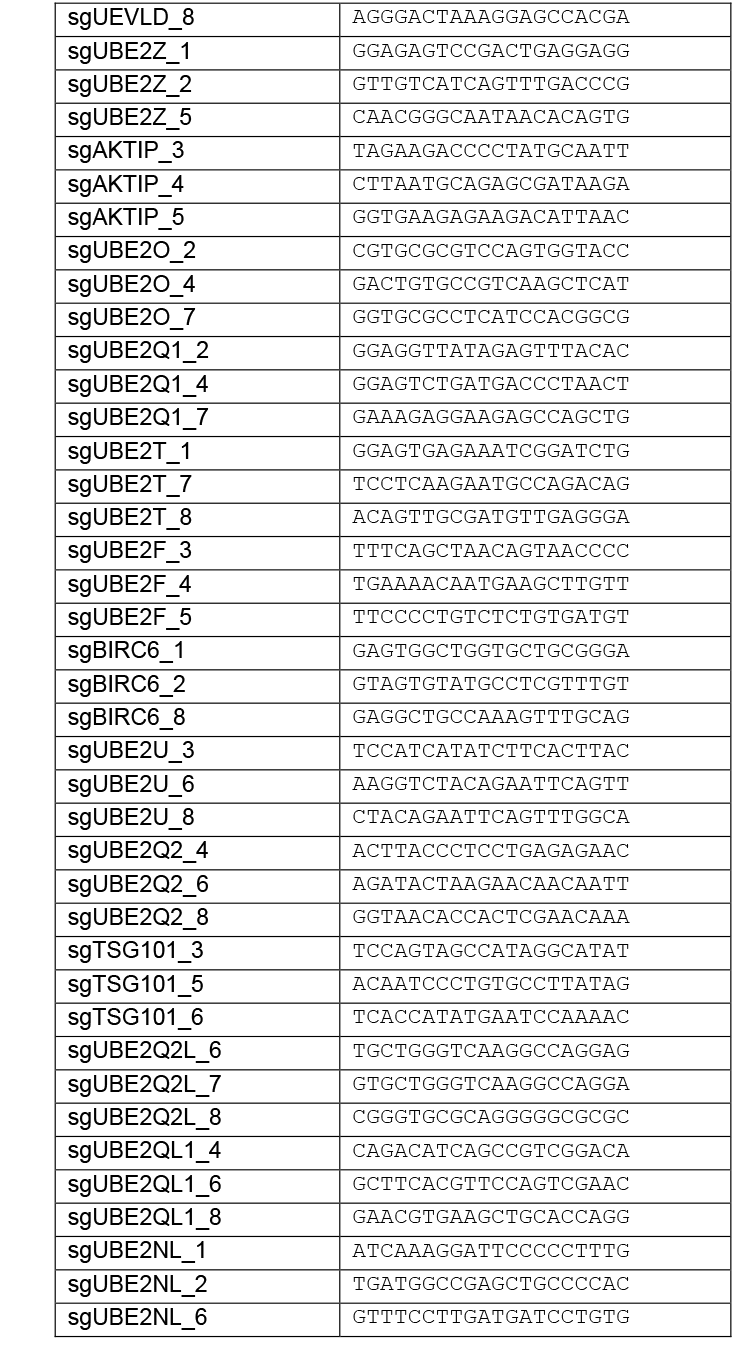
Guide RNA sequences targeting 41 annotated ubiquitin-conjugating enzymes.

